# Epigenomes in thermophilic microbial communities and their impact on the interaction across prokaryotes and mobilomes

**DOI:** 10.64898/2026.06.24.734251

**Authors:** Satoshi Hiraoka, Shigeru Shimamura, Keiko Usui, Yi Zhang, Tomomi Sumida, Yuya Tsukamoto, Shigeru Kawai, Arisa Nishihara

## Abstract

DNA chemical modifications, including methylation, are widespread in prokaryotes and mobilomes, including viruses, plasmids, and other extrachromosomal DNAs, and play important roles in their ecology and interactions. However, current knowledge of these modification systems and their association with interactions between hosts and mobilomes across communities, including those in extreme environments, is severely limited. Here, using single-molecule real-time sequencing and single-cell genome sequencing technologies, we conducted a culture-independent ‘metaepigenomic’ analysis of microbial communities in hot spring biofilms. A total of 248, 332, and 465 genomes were constructed from diverse prokaryotes, viruses, and extrachromosomal circular DNAs, respectively, from 10 biofilm samples collected from 3 hot spring sites. In total, 1106 candidate methylated motifs and 3280 genes associated with the restriction–modification (RM) system, including DNA methyltransferases (MTases), were identified. In contrast to the varied methylated motifs, the nucleotide-level modification ratios were consistent with those of a common *Escherichia coli* genome, and an environment-dependent epigenomic preference attributed to the lack of C5-methylcytosine was observed, as supported by direct measurements of modified bases by liquid chromatography–tandem mass spectrometry. A systematic survey revealed various defense systems in the genome, and almost half of the MTase genes were estimated to be genetically involved in defense mechanisms against extracellular DNA, such as RM systems. The mobilomes and their predicted hosts shared epigenomic patterns within each interactive subnetwork, suggesting that mobilome DNA was modified by host MTase during the current infection or transfection, rather than serving as historical records. Our findings highlight that DNA modification shapes multiple ecological and evolutionary strategies in interactions between prokaryotes and mobilomes, and that epigenomes serve as a potential signature for accurate prediction of current host–phage interactions.

## INTRODUCTION

Chemical modifications of DNA are found in diverse prokaryotes, viruses, and eukaryotes. DNA methylation is a representative DNA modification catalyzed by DNA methyltransferases (MTases), in which S-adenosylmethionine (SAM) provides the methyl group^1^. In prokaryotes, three methylation types (i.e.,N6-methyladenine [m6A], C5-methylcytosine [m5C], and N4-methylcytosine [m4C]) have been investigated in detail^2^. DNA methylation plays a role in regulating gene expression and DNA mismatch repair^3–5^. These systems perform various physiological functions, including asymmetric cell division^6,7^, ultraviolet (UV) tolerance^8^, motility^9^, virulence^10–12^, and adaptation to environmental changes under epigenetic control^13^. DNA methylation also facilitates cell protection from extracellular DNA (e.g., viral infection and plasmid transfection), known as restriction–modification (RM) systems^14,15^. The RM systems are classified into four types based on their subunit compositions and cofactor requirements. Types I, II, and III are composed of both MTase and restriction endonucleases (REase) and specify non-methylated DNA in most cases, whereas Type IV consists of only REase and specifies modified DNA substrates^16^. Some viruses possess MTases and modify their genomic DNA to escape the host Type I, II, and III RM systems. In contrast, Type IV systems have evolved to counter the viral anti-RM system, resulting in a co-evolutionary phage–host arms race^2,17^. Moreover, evidence of the horizontal gene transfer of RM system genes within and between domains and changes in MTase motif specificity have been frequently noted in the evolution of prokaryotes^18,19^. RM systems encoded on plasmids also facilitate host defense against phage infection, in association with methylation-mediated transcription regulation and their copy number^20,21^, and the RM systems play a co-evolutionary interaction between host and plasmids^22^. There has been increasing interest in exploring the various epigenomic systems amongst diverse prokaryotes and mobile genetic elements, or ‘mobilomes’ (e.g., viruses and extrachromosomal genomes such as plasmids) owing to their importance in microbial physiology, genetics, evolution, and disease pathogenicity^23–25^. However, most studies have relied on a small number of culturable prokaryotic strains; whereas, the majority of microbes remain uncultured, and the epigenomes in mobilomes have been overlooked overall. This small sample size skews our understanding of microbial epigenomics, particularly regarding its diversity, distribution, and effect on biological interactions.

Recent technological advances have led to the development of single-molecule real-time (SMRT) sequencing technology as a useful method for detecting DNA modifications. Its implementation on PacBio sequencing platforms has yielded an array of DNA modifications among prokaryotic^26–31^ and viral strains^32,33^. Based on this innovative technique, a culture-independent shotgun metagenomic and epigenomic analytical approach referred to as metaepigenomics has been developed (Figure 1). Using this methodology, community-level epigenomic analyses have been conducted on several environmental samples (e.g., freshwater and seawater), demonstrating diverse DNA modifications in prokaryotes^34–37^ and viruses^34,38^ that may be associated with microbial ecology and evolution.

**Figure 1.**
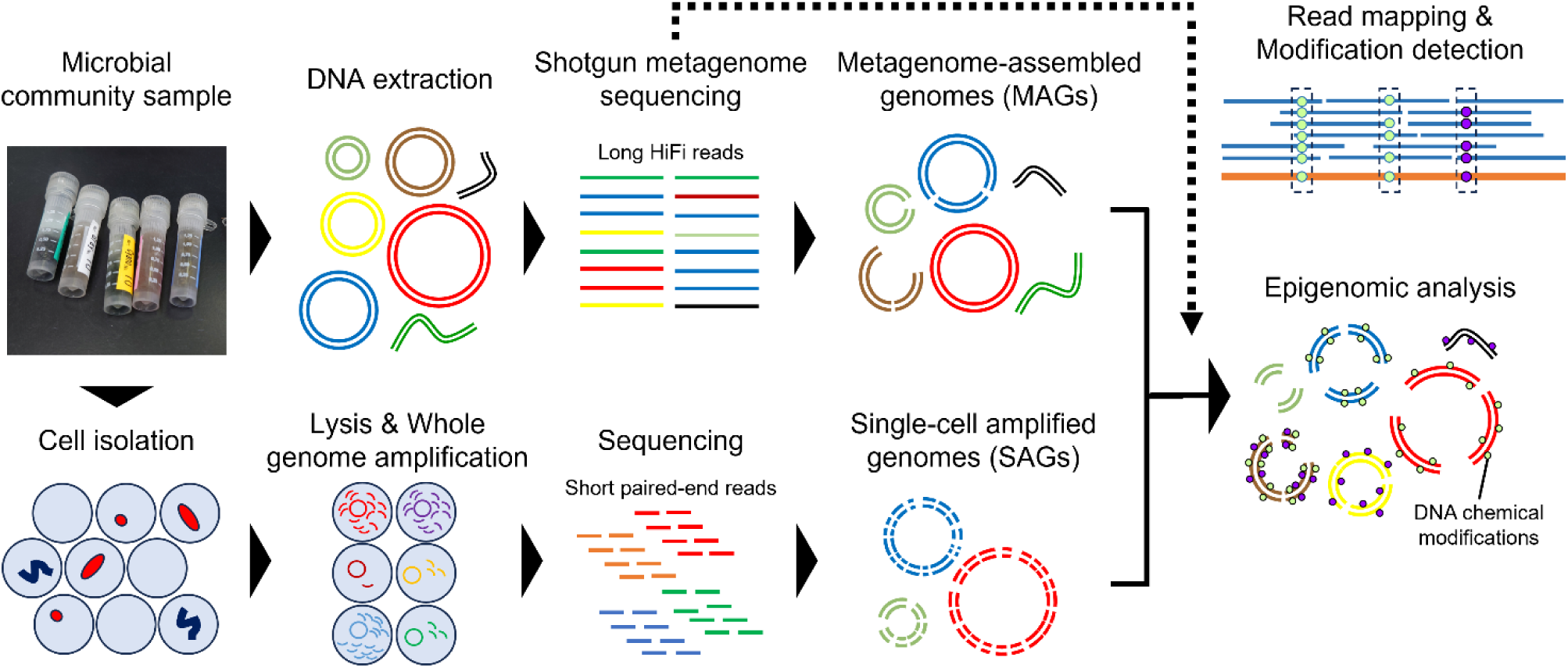
Schematic view of metaepigenomics in the present study.

Here, we conducted a meta-epigenomic analysis of hot spring microbial communities to reveal the epigenomes of diverse thermophilic prokaryotes and mobilomes. Hot spring biofilms establish semi-closed, high-biomass, thermophilic communities shaped by water chemistry and spatial temperature gradients, the epigenomic status of which remains largely unknown. In addition, abundant mobilomes play significant roles in hot spring ecosystems^39^, although they remain poorly investigated. Thus, hot spring biofilms are an ideal system for examining the epigenomic diversity across taxa and the associations that underlie the interactions between prokaryotes and mobilomes. Diverse DNA modifications were successfully characterized in numerous metagenome-assembled genomes (MAGs) and single-cell amplified genomes (SAGs) across prokaryotes and mobilomes. A portion of the prokaryotic epigenomic patterns was associated with phylogeny rather than habitat, suggesting vertical propagation, whereas the sporadic occurrence of methylated motifs in other clades may reflect horizontal transfer or motif diversification. Epigenomic patterns were similar between mobilomes and host prokaryotes, demonstrating that the acquisition of methylation is mediated by host MTases during infection or transfection. In particular, the conservation or diversification of motifs within each clade is associated with the size of the host–mobilome interaction network, suggesting two distinct ecological epigenomic roles, which can be referred to as “exchange” and “retention” strategies.

## MATERIAL AND METHODS

### Hot spring biofilm sampling and water chemical analysis

Biofilm samples were collected at three terrestrial hot springs in Japan: two sites at Nakabusa (the stream site *Kassennoyu* [36° 23.55N, 137° 44.88E] and the wall site *Kojikinoyu* [36° 23.33N, 137° 44.88E]), one site at Shiriyaki (36° 38.89N, 138° 38.46E), and one site at Dewanoyu (40° 25.71N, 140° 37.23E) from June, 2021 to October, 2022 (Note S1; Figure S1; Tables S1 and S2). Nakabusa is a sulfidic, slightly alkaline hot spring, and has been intensively studied with regard to its microbial diversity and function^40,41^. In the Nakabusa hot spring, six biofilm samples (NSBL, NSGRY, NSOL, NSGRN, NSGF, and NSBR) were collected at a temperature range from 86 °C to 51 °C at the stream site along with the artificial stream line, and two samples (NWPT and NWRS) were collected at 75 °C and 66 °C at the wall site where hot spring water emerged from cracks on the erosion control dam. Two samples were collected at the source of Shiriyaki and Dewanoyu hot springs at 51 °C and 58 °C, respectively (referred to as SY and DW, respectively). Shiriyaki is a slightly acidic, sulfate-rich, moderate-temperature hot spring and, to the best of our knowledge, its native microbial communities have not yet been reported on. Dewanoyu is a sodium chloride-rich, moderately thermophilic hot spring with a neutral pH in which microbial communities have not been investigated to the best of our knowledge. Sampling was permitted by Nakabusaonsen at the Nakabusa hot spring sites, whereas the other sites were exempt from specific sampling permits. The samples were collected using sterile tweezers and placed in sterile 2.0-mL screw-cap tubes, stored in a cool box on dry ice, and immediately transferred to the laboratory where they were stored at temperatures below −30 °C.

For chemical measurements, except for sulfide, hot spring water was filtered (0.22 µm) into sterile 100-mL Falcon tubes. The total Fe and Mn concentrations were measured using an Agilent 5100 VDV ICP-OES (Agilent Technologies, Santa Clara, California, USA). The anion and cation concentrations were determined by ion chromatography (Shimadzu, Kyoto, Japan) equipped with a conductivity detector (CDD-10A series), Shim-pack IC-SA2 column for anion analysis, and Shim-pack IC-C4 column for cation analysis. For sulfide analysis, 2 mL of hot spring water was filtered (0.22 µm) and fixed with 500 µL of 90 µM zinc acetate solution in N₂-purged 7-mL glass vials. The sulfide concentration was determined colorimetrically using the methylene blue method^42^. The collected samples were stored in a cool box on ice and transferred to the laboratory where they were then stored at 4 °C prior to analysis.

### Metagenomic sequencing and assembly

Microbial DNA for sequencing was retrieved using a DNeasy PowerSoil Pro Kit (QIAGEN, Hilden, Germany) according to the supplier’s protocol. SMRT sequencing was conducted using a PacBio Sequel II system (Pacific Biosciences of California, Menlo Park, California, USA) at Macrogen (Seoul, Korea). High-fidelity (HiFi) reads with > 99% average base-call accuracy were retained using the standard PacBio SMRT software package with default settings. Metagenomic HiFi read coverage was estimated using Nonpareil3 with default settings^43^. For taxonomic assignment of HiFi reads, Kaiju^44^ in Greedy-5 mode (‘-a greedy -e 5’ settings) with refseq_nr (NCBI RefSeq non-redundant protein) prebuild database (date: 2024-08-13) was used. Coding sequences (CDSs) were predicted using Prodigal^45^ in anonymous mode (‘-p meta’ setting).

The HiFi reads were used in a two-step *de novo* assembly workflow (Figure S2). First, the reads were subjected to a metagenomic HiFi read assembler, hifiasm-meta^46^ with the default settings. Second, to resolve bins with complex assembled graphs, HiFi reads were mapped to assembled graphs comprising > 3 contigs and >100 kb in total length using pbmm2 (https://github.com/PacificBiosciences/pbmm2) with the default setting. The mapped reads were then retrieved and used for the re-assembly using hifiasm^47^, which was designed for HiFi reads from isolated organisms, with ‘-l0’ setting. The assembled contigs from both the first and second steps were evaluated for topology, continuity, and quality. Briefly, all circular contigs and linear contigs with a length > 500 kb were used for genome quality assessment using CheckM2^48^, and those with medium (> 50% completeness and < 10% contamination) or high (> 90% completeness and < 5% contamination) quality were identified as prokaryotic MAGs (pMAGs). Quality thresholds were based on the minimum information about a metagenome-assembled genome (MIMAG) standards^49^.

For viral sequence collection, both the first- and second-assembled contigs were screened using VirSorter2^50^ with the default settings. Quality assessment and removal of flanking host regions from the integrated proviruses were performed using CheckV^51^. Sequences assigned to either ‘Complete,’ ‘High-quality,’ or ‘Medium-quality’ were retrieved according to the minimum information about an uncultivated virus genome (MIUViG) ^52^. The sequences after dereplication using Galah (https://github.com/wwood/galah) with ‘--ani 95’ setting (genomes with average nucleotide identity [ANI] > 0.95 similarity will be dereplicated) were defined as viral MAGs (vMAGs).

For the collection of extrachromosomal sequences such as plasmids, both first- and second-assembled contigs were examined. Notably, considering the technical challenges and aiming for precise analysis, we focused on the candidate complete extrachromosomal genomes with a circular structure in this study: among the contigs, those with a circular structure and < 10% completeness estimated by CheckM2^48^ were retrieved. The contigs after dereplication using Galah with ‘--ani 95’ setting were defined as extrachromosomal circular MAGs (eccMAGs).

### Single-cell genome sequencing analysis

To obtain genomes that were difficult to reconstruct from metagenomic data alone, we performed single-cell genome sequencing on the NSOL sample. Library preparation, sequencing, and assembly were performed at bitBiome (Tokyo, Japan). Briefly, microbial cells were isolated from the samples and sheared onto gel beads using the bitBiome microbiome analysis platform (bitMAP)^53,54^. After cell lysis and subsequent whole-genome amplification, only fluorescence-positive gel beads stained with 10× SYBR Green I (Thermo Fisher Scientific) were collected. The retrieved amplified DNAs were subjected to library preparation using xGen DNA Lib Prep EZ UNI (Integrated DNA Technologies, Coralville, Iowa, USA) and sequenced on a DNBSEQ-G400 (MGI Tech, Shenzhen, China) with 2 × 150 bp settings. Adopters were removed using the BBDuk script implemented in BBtools (https://github.com/BioInfoTools/BBMap) with ‘ktrim=r ref=adapters k=23 mink=11 hdist=1 tpe tbo qtrim=r rimq=10 minlength=40 maxns=1 minavgquality=15’ settings. Contaminated sequences derived from humans were removed using bbmap with ‘quickmatch fast untrim minid=0.95 maxindel=3 bwr=0.16 bw=12 minhits=2 path=human_masked_index(*) qtrim=rl trimq=10’ settings. The remaining reads were assembled using SPAdes^55^ with ‘--sc --careful --disable-rr --disable-gzip-output –k 21,33,55,77,99,127’ settings. Contigs < 200 bp in length were removed using SeqKit2^56^. The assembled genomes of medium (> 50% completeness and < 10% contamination) and high quality (> 90% completeness and < 5% contamination) were defined as prokaryotic SAGs (pSAGs), according to the minimum information about a single amplified genome (MISAG) ^49^.

### Bioinformatic analyses of HiFi reads and genomes

The pMAGs and pSAGs were dereplicated using dRep^57^ with ‘-comp 0 -con 0 --primary_chunksize 500 -sa 0.99 -l 50000’ settings (genomes with ANI > 0.99 similarity were dereplicated, anticipating strain-level representatives, and those with a length < 50 kb were removed) and were defined as pMAG/SAGs. The taxonomy was estimated using the GTDB-Tk with r226 database^58^. Genome annotation was partially performed using DFAST^59^. CDSs were predicted using Prodigal^45^ with the default settings. For the total phylogenetic analysis of prokaryotic genomes, a maximum-likelihood tree was constructed using PhyloPhlan3^60^ on the basis of a set of 400 conserved prokaryotic marker genes^61^ with ‘--force_nucleotides --diversity high --accurate’ settings. To calculate genome coverage, HiFi reads were mapped onto the genomes using pbmm2 with default settings. Short sequence repeats (SSRs) were detected using a Tandem repeats finder (TRF) ^62^. Mapped reads were visualized using Integrative Genomics Viewer^63^.

For the vMAGs, taxonomy was estimated using VITAP^64^ with the official prebuild database DB_VMR-MSL_v40. CDSs were predicted using Prodigal^45^ in an anonymous mode (‘-p meta’ setting). A proteomic tree of vMAGs was constructed using ViPTreeGen^65^ with default settings. Functional annotations and genome coverage calculations were performed in the same manner as for the pMAG/SAGs.

For the eccMAGs, CDSs were predicted using Prodigal^45^ in an anonymous mode (‘-p meta’ setting). Functional annotations and genome coverage calculations were performed in the same manner as for pMAG/SAGs. Plasmids were predicted using Plasmer^66^. To construct a dendrogram based on genome similarity distance, ANI was calculated using fastANI with ‘--minFraction 0.05 --fragLen 1000’ settings. The calculated distance matrix was then subjected to hierarchical clustering using the Ward’s method.

Genes encoding DNA MTases, REases, and DNA sequence recognition proteins (S subunits) were searched using DIAMOND^67^. The retrieved RM system genes were compared against an experimentally confirmed gold-standard dataset from REBASE^68^ (last updated on December 28, 2025), with a cutoff e-value of ≤ 1E-5. Information including sequence specificity and protein type for each RM system gene was retrieved from the REBASE gold standard dataset. Defense systems were predicted using PADLOC with the official prebuild database PADLOC-DB v2.0.0^69^. For reliable investigation, the systems yet validly classified (’XXX_other’) were excluded. Host prediction of vMAGs and eccMAGs was conducted using iPHoP^70^ with an in-house host genome database composed of the official database iPHoP_db_Jun25_rw in addition to the pMAG/SAGs reconstructed in this study.

### Epigenomic analysis

DNA modification detection and motif analyses were performed using SMRT Link (v25.0). To avoid multiple mapped reads, HiFi reads were first mapped to the full set of genomes with pbmm2, and the mapped reads for each genome were retrieved and used for modification detection and motif prediction after individual remapping using the same method. Candidate motifs with scores above the default threshold values were retrieved as the modified motifs. Motifs with several ambiguous sequences which were considered to have occurred because of misdetection were manually curated. For example, TTCGA**A**NNNNNNNNNNNNNG was detected in pMAG_1st_SY_009, where N = A/T/C/G; however, this motif represents palindromic <u>T</u>TCGA**A**. Furthermore, the spurious partial sequences of the latter NNNNNNNNNNNNNG were likely due to incomplete detection of the motif. We frequently found candidate motifs exhibiting this type of ambiguity in vMAGs and eccMAGs, consistent with our previous observations^34^. This may have resulted from the limited capacity of the method to estimate motifs in extremely small genomes, whereby the low abundance of motif sequences in these genomes adversely affected the motif-finding algorithm implemented in MotifMaker, which is part of the SMRT Link software package. For comparative epigenomic analysis, the gold-standard HiFi reads of the *E. coli* K-12 MG1655 genome generated using the PacBio Sequel II platform were downloaded (https://downloads.pacbcloud.com/public/dataset/Ecoli/egs/) and used.

### Nucleosides analysis

To estimate the DNA modification ratio in the metagenome, modified and unmodified nucleosides were directly quantified using liquid chromatography–tandem mass spectrometry (LC-MS/MS). Metagenomic DNA from hot spring biofilm samples was extracted using the same method as that used for the sequencing analysis. For comparison, we also used genomic DNA from *Escherichia coli* DH5α (K-12 derivative), which harbors three active MTase genes: M.EcoKI (A**A**CNNNNNNG<u>T</u>GC/GC**A**CNNNNNNG<u>T</u>T specificity), Dam (G**A**<u>T</u>C), and Dcm (C**C**W<u>G</u>G, m5C). The *E. coli* DH5α competent cells (Takara Bio, Shiga, Japan) were cultured in Luria–Bertani (LB) broth, and genomic DNA was extracted using a DNeasy UltraClean Microbial Kit (QIAGEN) according to the manufacturer’s protocol. DNA (500 ng per sample) was enzymatically digested at 37 °C for 1 h into nucleosides using Nucleoside Digestion Mix (New England Biolabs [NEB], Ipswich, Massachusetts, USA) according to the supplier’s protocol. The resulting digest was directly subjected to nucleosides analysis with LC-MS/MS.

LC-MS/MS was performed using a previously described protocol, with slight modifications^71^. The LC-MS/MS system consisted of an ultra-high-performance liquid chromatography pump (UltiMate 3000, HPG-3200RS, Thermo Fisher Scientific) and an Orbitrap Fusion mass spectrometer (Thermo Fisher Scientific). The LC was performed on a separation column (Acclaim 120 C18: 2.1 mm of diameter, 150 mm of length, 2.2 μm of particle size; Thermo Fisher Scientific) with a flow rate of 0.3 mL/min at 35 °C. Solvent A (0.1% formic acid in water) and Solvent B (0.1% formic acid in methanol) were used as the mobile phases. A gradient of 0–3 min 5–15% B, 3–5 min 20% B, and 5–8 min 5% B was used. MS analysis was operated in positive electrospray ionization mode to detect targeted nucleosides on their accurate masses. The full scan MS settings included: Detector Orbitrap, Resolution 120,000, Quadrupole Isolation ON, Scan Range 100–450 *m/z*, an acquisition gain control (AGC) Target Standard, Maximum Injection Time Mode Auto, S-lens Radio Frequency Level 60%, EASY-IC ON, Spray Voltage 3500 V, Sheath Gas Flow Rate 50 arbitrary units, Aux Gas Flow Rate 10 arbitrary units, Ion Transfer Tube Temperature at 250 °C, and Vaporizer Temperature at 220 °C. The obtained data were analyzed using Qual Browser in Xcalibur (version 4.3.73.11) (Thermo Fisher Scientific). Regarding the standard substances: 2′-deoxyadenosine (dA), 2′-deoxycytidine (dC), and 2′-deoxy-5-methylcytosine (m5dC) were purchased from Tokyo Chemical Industry (Tokyo, Japan); 2′-deoxy-N6-methyladenine (m6dA) was purchased from Santa Cruz Biotechnology (Dallas, Texas, USA); 2′-deoxy-N4-methylcytidine (m4dC) was purchased from BOC Sciences (Shirley, New York, USA).

## RESULTS

### Community structure and MTase gene distribution in hot spring biofilm communities

Ten biofilm samples were collected from three hot springs (Nakabusa, Shiriyaki, and Dewanoyu) in Japan (Figure S1; Table S1). The three hot springs are geographically distant and have different water chemical characteristics and temperature (51–86 ℃) (Note S1; Table S2). For each sample, PacBio Sequel II produced 0.44–3.0 million (5.5–31.1 Gb) HiFi reads with > 99% accuracy, with the average length ranging from 7667–12934 bp (Figure S3; Table S3). The HiFi reads were estimated to cover 90–97% of the community diversity per sample, indicating that the sequencing data covered almost the entire metagenome (Figure S4). The taxonomic assignment of the HiFi reads showed that the prokaryotic community structures differed markedly between samples (Figures 2A–C; Note S2). In samples from Nakabusa hot spring, where living thermophilic microbes have been studied for decades, the community composition was consistent with that of previous studies based on 16S rRNA amplicon sequencing^40,72^, suggesting low temporal fluctuations in the community structure at the sampling sites and minimal sequencing biases in the analysis.

**Figure 2.**
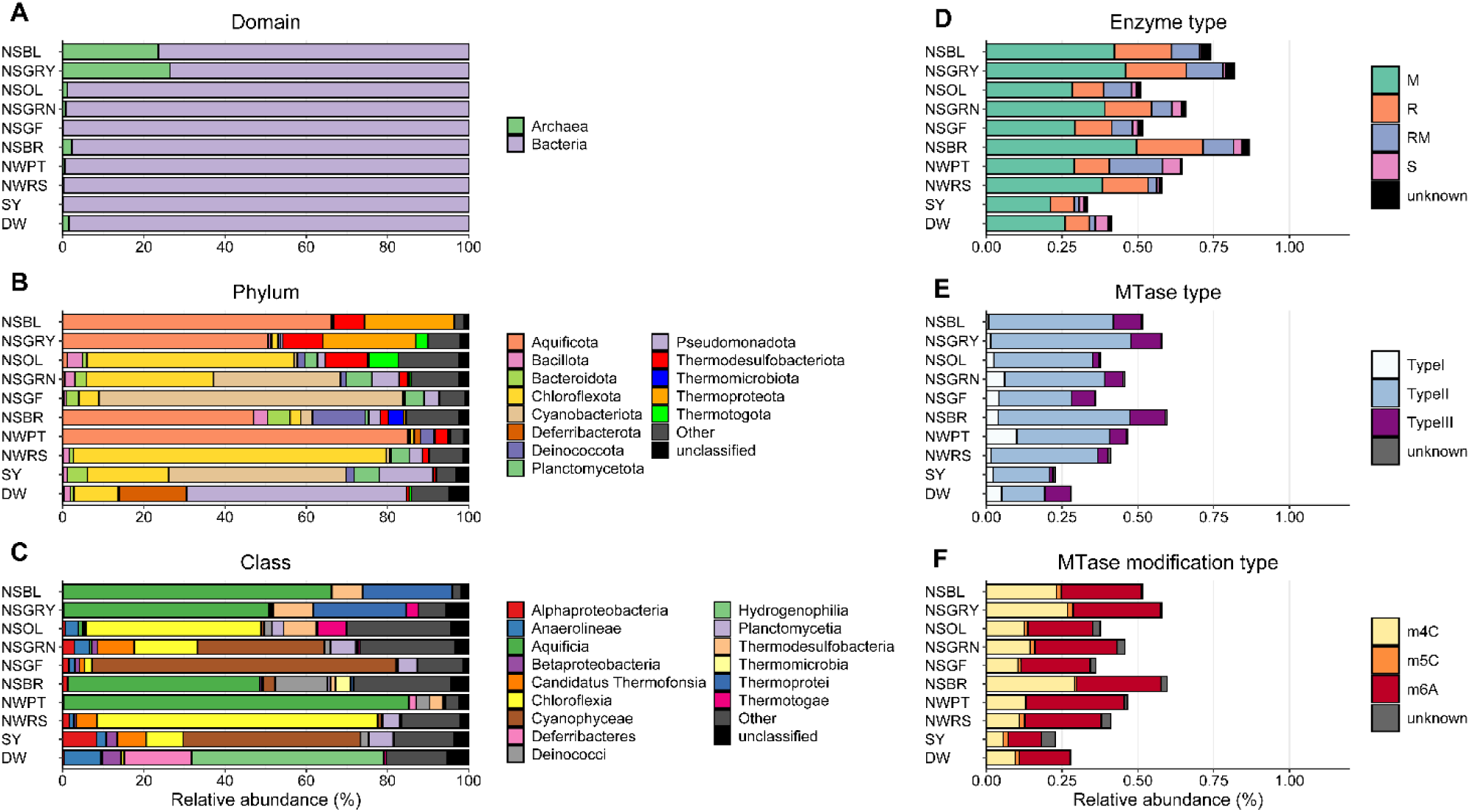
Profiles of taxonomy and RM system genes in hot spring metagenomes. (**A–C**) Taxonomic profiles of metagenomic HiFi reads. The estimated relative abundances obtained using Kaiju with the NCBI RefSeq non-redundant protein database at the (**A**) domain, (**B**) phylum, and (**C**) class levels are presented. Sequences assigned to members with < 3% abundance across the samples were grouped as Others. (**D**–**F**) Profiles of genes encoding DNA restriction and modification enzymes per total CDSs on HiFi reads: (**D**) distribution of enzyme types: DNA methyltransferase (MTase; M), restriction endonuclease (REase; R), protein fused with M and R domains (RM), and DNA sequence-recognition protein (S); (**E**) distribution of MTase types; (**E**) distribution of modification types.

Genes related to DNA methylation and RM systems were determined by systematic annotation of MTase and REase genes on the HiFi reads using the REBASE gold standard database^68^. In general, genes assigned to MTase (M), REase (R), a protein fused with the MTase and REase domains (RM), and a DNA sequence-recognition protein involved in Type I RM systems (S) showed different total abundances but similar compositions across the microbial communities (Figure 2D). Within the MTase proteins (i.e., M and RM), Type II was predominant, accounting for 50.9–86.9% (74.2% on average) of each sample (Figure 2E). The relative abundances of Types I (1.5–18.2%, 9.6% on average) and III (5.3–30.8%, 15.0 on average) were approximately 3-times lower and 2-times higher, respectively, than those identified in the genomes of wide prokaryotic isolates (27% and 8%, respectively)^73^. Among the detected MTases, the most abundant predicted modification type was m6A (46.7–63.2%, 56.6% on average), followed by m4C (24.2–48.8%, 34.7% on average) (Figure 2F). Notably, the other m5C MTases were almost scarce throughout the samples (0.4–7.5%, 3.4% on average). Overall, the composition of MTases and modification types were similar across samples, even among chemically distinct hot spring sites and different community structures.

### Diverse epigenomes in hot spring biofilm communities

From metagenome and single-cell genome sequencing analyses, we obtained 248 prokaryotic, 332 viral, and 465 extrachromosomal circular genomes (referred to as pMAG/SAGs, vMAGs, and eccMAGs, respectively) (Note S3; Figures S5 and S6; Table S4; Data S1). Among the prokaryotic genomes, 117 were predicted to be of high quality, and 85 were circular, demonstrating the high performance of metagenomic assembly using long, highly accurate reads. The genomes showed a highly localized abundance distribution among the samples, indicating isolated semi-closed communities at each sampling station across all three genome types (Figure S7).

From the pMAG/SAGs, a total of 1138 candidate modified motifs were detected from 157 (63%) genomes (Data S2). From the vMAGs and eccMAGs, 923 and 2582 tentative motifs were detected, respectively, although many of these were ambiguous owing to the low detection power of the modified motifs from small genomes, as we discussed previously^34^. Therefore, we considered only the motif sets predicted from pMAG/SAGs in the following analysis. The motifs comprised 353 unique motifs, including 91 palindromic motifs that allowed for double-strand modifications. Among these, 163 and 164 were classified as m6A and m4C methylation, respectively. Although current SMRT sequencing technology does not support m5C detection, we identified 25 candidate m5C motifs. The modification types of the other motifs were unclassified. Among the motifs, G**A**<u>T</u>C was detected most frequently (63 pMAG/SAG), followed by G**A**N<u>T</u>C (56 pMAG/SAG), **C**CG<u>G</u> (53 pMAG/SAG), **C**GC<u>G</u> (37 pMAG/SAGs), and G<u>T</u>**A**C (33 pMAG/SAGs), where N = A/C/G/T, and the bold and underlined characters indicate modified bases and those on the complementary strand, respectively. Notably, even considering some vague motifs, 264 motifs did not match the known MTase motifs in the REBASE repository, e.g., C**C**TGG(m4C), GTC**C**AGG(m4C), GGNN**C**CG(m4C), CAA**C**CG(m4C), CAA**C**CGA(m4C), CAA**C**T(m4C), GTC**C**T(m4C), CGA**A**CA, CGACG**A**, GA**C**TCA(m4C), GCC**A**G, C**C**TCC(m4C), GAG**C**CC(m4C), and GGG**C**TC(m4C). In addition, methylated motifs likely catalyzed by Type I MTases, which are generally characterized as bipartite sequences with a gap of unspecified nucleotides (e.g., **A**TGNNNNNTAC), were scarcely detected (155 motifs) across the pMAG/SAGs. This result was concordant with the low abundance of Type I MTases in the metagenome (Figure 2D), suggesting that Type I RM systems were rare in the hot spring communities.

To further validate the DNA modifications, we conducted an LC-MS/MS analysis of total nucleosides in each metagenome. The modification ratios of m6dA/dA were high across the hot spring samples (11.8% on average), followed bym4dC/dC (2.1%) (Figure S8). In sharp contrast, dm5C was largely absent in many samples, and the dm5C/dC ratio was extremely low across samples (0.1%). This was consistent with the meta-epigenomic analysis, which indicated that m6A and m4C motifs were abundant, whereas m5C motifs were poorly detected (Data S2). This was consistent with the functional annotation of the metagenomes, which showed that m6A- and m4C-type MTases were abundantly detected, whereas m5C-type MTases were scarce overall (Figure 2F). In contrast to the hot spring samples, measurements of *E. coli* DH5α genomic DNA showed high m6dA/dA and m5dC/dC ratios, and an absence of dm4C. This result exactly matched the expected profiles of *E. coli* DH5α, in which A**A**CNNNNNNG<u>T</u>GC/GC**A**CNNNNNNG<u>T</u>T (m6A), G**A**<u>T</u>C (m6A), and C**C**W<u>G</u>G (m5C) are methylated by native MTases M.EcoKI, Dam, and Dcm, respectively, and hence supported the validity of the direct modification measurements. Consequently, it can be concluded that the hot spring biofilm microbes generally lacked the ability to modify C to m5C.

We further investigated the base-level modification ratios in prokaryotic genomes using the SMRT sequencing data. Among the pMAG/SAGs, the modified base ratios were 0.40%, 0.31%, and 0.30% for m4C, m5C, and m6A, respectively. The ratios per genome were slightly fractionated but were similar across phyla (Figure S9A), and no obvious trends were observed between samples (Figure S9B). However, these values were questionable because m5C modifications were abundantly detected across genomes and samples, which was inconsistent with the motif prediction and nucleoside analysis using LC-MS/MS described above. Indeed, the modification ratios in the *E. coli* genome were 0.27%, 0.72%, and 0.45% for m4C, m5C, and m6A, respectively; whereas, the theoretical values, accounting for the three native MTases were 0.00%, 0.26%, and 0.41%, respectively. These results clearly indicated overdetection, particularly for m4C and m5C methylation, resulting from the low reliability modification detection power of the current SMRT sequencing technology at the nucleotide base level, as previously described^74^. Although there were biases, the modification ratios in hot spring microbial genomes and metagenomes were largely comparable to those in *E. coli* genomes, consistent with the nucleoside analysis. These trends suggested that the microbes maintained consistent levels of nucleotide methylation across lineages and habitats, possibly to avoid harmful biological effects such as gene expression disorders.

### Phylogenetic distribution of methylated motifs

To investigate the phylogenetic distribution of the DNA modification system, we visualized the modification ratios of detected methylated motifs across genomes. Within prokaryotes, the distribution of the motif methylation was largely associated with phylogenetic clades (Figure 3). For example, within the phylum Planctomycetota, <u>G</u>CG**C** were found in the genomes of all organisms. Within the phylum Aquificota, motifs such as **C**CG<u>G</u> and G**A**<u>T</u>C were distributed across genomes. Among the genomes of the Aquificaceae family in Aquificota, **C**GC<u>G</u>, C**C**W<u>G</u>G, **C**TA<u>G</u>, GG<u>G</u>**C**CC, G**A**N<u>T</u>C, and G<u>T</u>**A**C were widely detected. Within the phylum Desulfobacterota, G<u>T</u>**A**C was detected widely. In addition, **C**CG<u>G</u> and C**C**TC were found in the MAG clade; whereas, GC**A**NNNNNNNGTA, GC**A**NNNNNNNGTG, T**A**CNNNNNNNTGC, and C**A**CNNNNNNNTGC were found in the SAG clade. Within the phylum Armatimonadota, many motifs (e.g., **C**TA<u>G</u>, C**C**W<u>G</u>G, <u>G</u>GNC**C**, A<u>G</u>**C**T, G**A**N<u>T</u>C, C**A**<u>T</u>G, C<u>T</u>CG**A**G, and GC**A**<u>T</u>GC) were spread in the genomes from the class Fimbriimonadia, but were not detected in organisms from the other classes. These results demonstrated an association between the phylogeny and epigenome. In sharp contrast, some lineages showed a sporadic distribution of motifs with no clear association with the phylogenetic topology. For example, in Chloroflexota, Bacteroidota, and Spirochaetota, many motifs were detected in different combinations across genomes. These clades tended to harbor more MTase genes than the former motif-conserved clades (11.5 and 6.3 genes, respectively, on average) (p < 0.05, t-test). In the case of viral genomes, further sparse and sporadic distributions were observed overall (Figure S10). Extrachromosomal circular genomes also showed a sporadic distribution overall, but highly conserved motifs within specific clades were observed (Figure S11).

**Figure 3.**
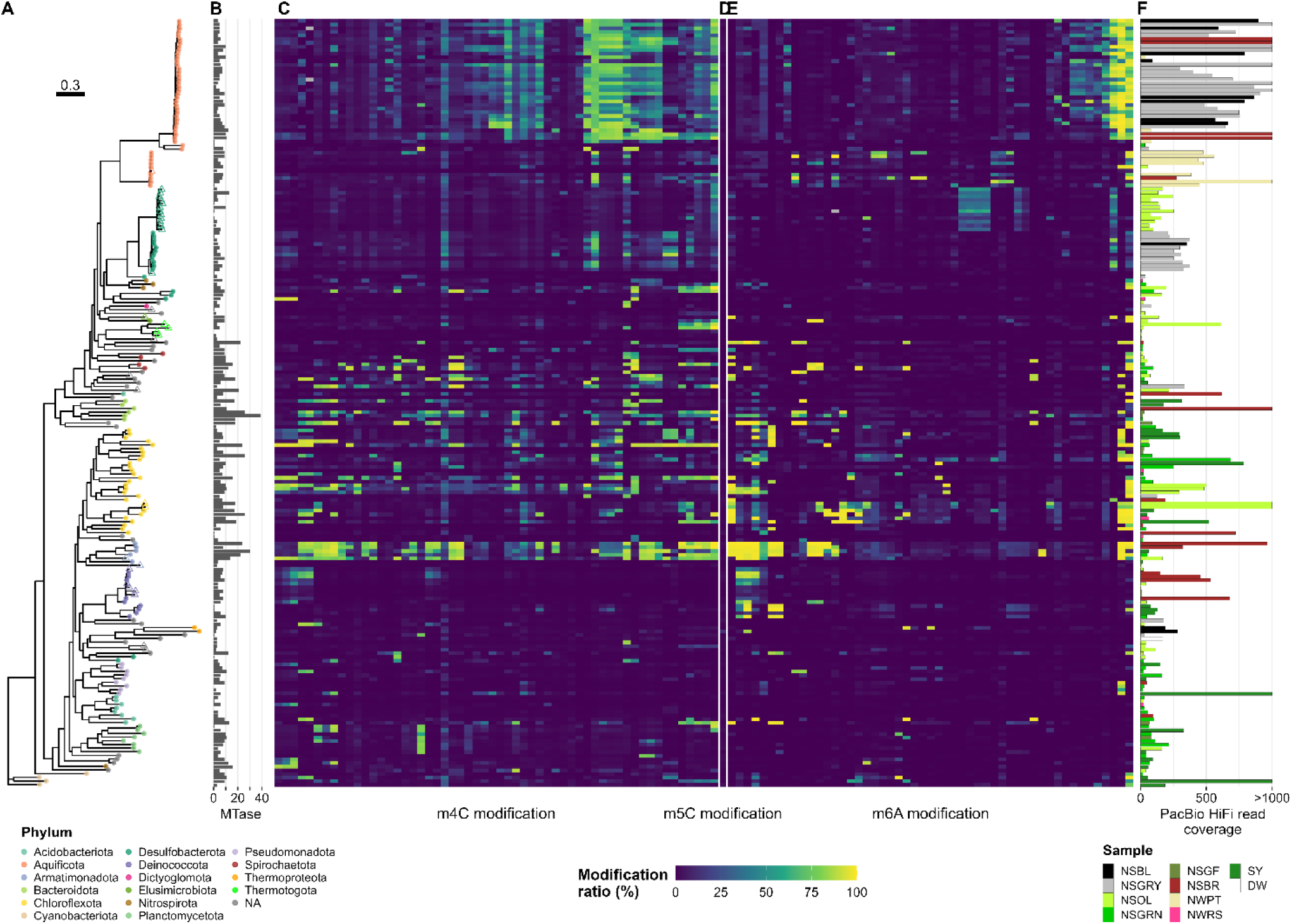
Methylomes of prokaryotic genomes. (**A**) A phylogenetic tree was constructed based on a set of up to 400 conserved bacterial marker genes using the maximum-likelihood method. Nodes indicated by a filled circle and an open triangle represent a metagenome-assembled genome (MAG) and a single-cell amplified genome (SAG), respectively. Node color indicates taxonomy at the phylum level. (**B**) Number of MTase genes identified in each genome. (**C**–**E**) Modification ratios of detected motifs per genome: (**C**) m4C; (**D**) m5C; and (**E**) m6A. Motifs detected in at least two prokaryotic genomes, excluding spurious sequences, were used. The color range from blue to green to yellow represents the modification ratios of motifs across genomes. The underlined characters indicate modified bases. Notably, modification ratios were affected by overlapped motif sequences; for example, GCG**C**TC was completely overlapped by <u>G</u>CG**C**, and both motifs showed similar modification rates in their genomes, except one genome (pMAG_1st_NSGRN_Circle_005), where GCG**C**TC was detected on the genome as per the metaepigenomic analysis, and concordantly, the modification ratio of GCG**C**TC was higher than that of <u>G</u>CG**C**. Motifs were ordered by hierarchical clustering of modification ratios using Euclidean distance and the Ward’s method. (**F**) Coverage of HiFi reads on each genome. The bar color represents the source of the genome and HiFi reads.

Across all the prokaryotic genomes in this study, no obvious clades were absent from the methylated motif. This trend was distinct from that of our previous study on marine microbial communities, in which many clades, such as Gemmatimonadota (formerly known as Gemmatimonadetes), a part of Pseudomonadota (Gammaproteobacteria), Nitrospinota (Nitrospinae), Verrucomicrobiota (Verrucomicrobia), and Myxococcota (Deltaproteobacteria), showed no methylated motifs on their genome^34^. This may have reflected differences in immune strategies between habitats.

### Defense systems and epigenomic systems in reconstructed genomes

We investigated the prevalence of defense systems, including those mediated by DNA modifications (e.g., RM systems), in communities to examine potential interactions between prokaryotes and mobilomes. The pMAG/SAGs harbored an average of 8.2 defense systems (≤ 27) across their genomes (Figure 4A; Data S1). CRISPR-Cas and RM systems were the two most abundant and frequently observed defense systems across the pMAG/SAGs (average of 1.8 and 1.6 systems per genome, respectively), with the exception of systems that are currently not well verified, such as phage defense candidates (PDC)^69^ and PD^75^ systems (Figures 4B and C). The RM system types were similar across lineages at the phylum level, whereas the distribution of the CRISPR-Cas system subtypes showed an association (Figure S12). Among the four most abundant systems, no clear relationship was observed in the number of systems per genome (Figure S13). Other known antiviral defense systems associated with DNA modification, such as BREX^76^ and DISARM^77^ were also predicted despite their low frequency. Among the vMAGs, a few defense systems were detected, including eight RM and eight CRISPR-Cas systems. In contrast to viral genomes, comparatively larger numbers of defense systems were detected on eccMAGs, including 51 RM and 12 CRISPR-Cas systems.

**Figure 4.**
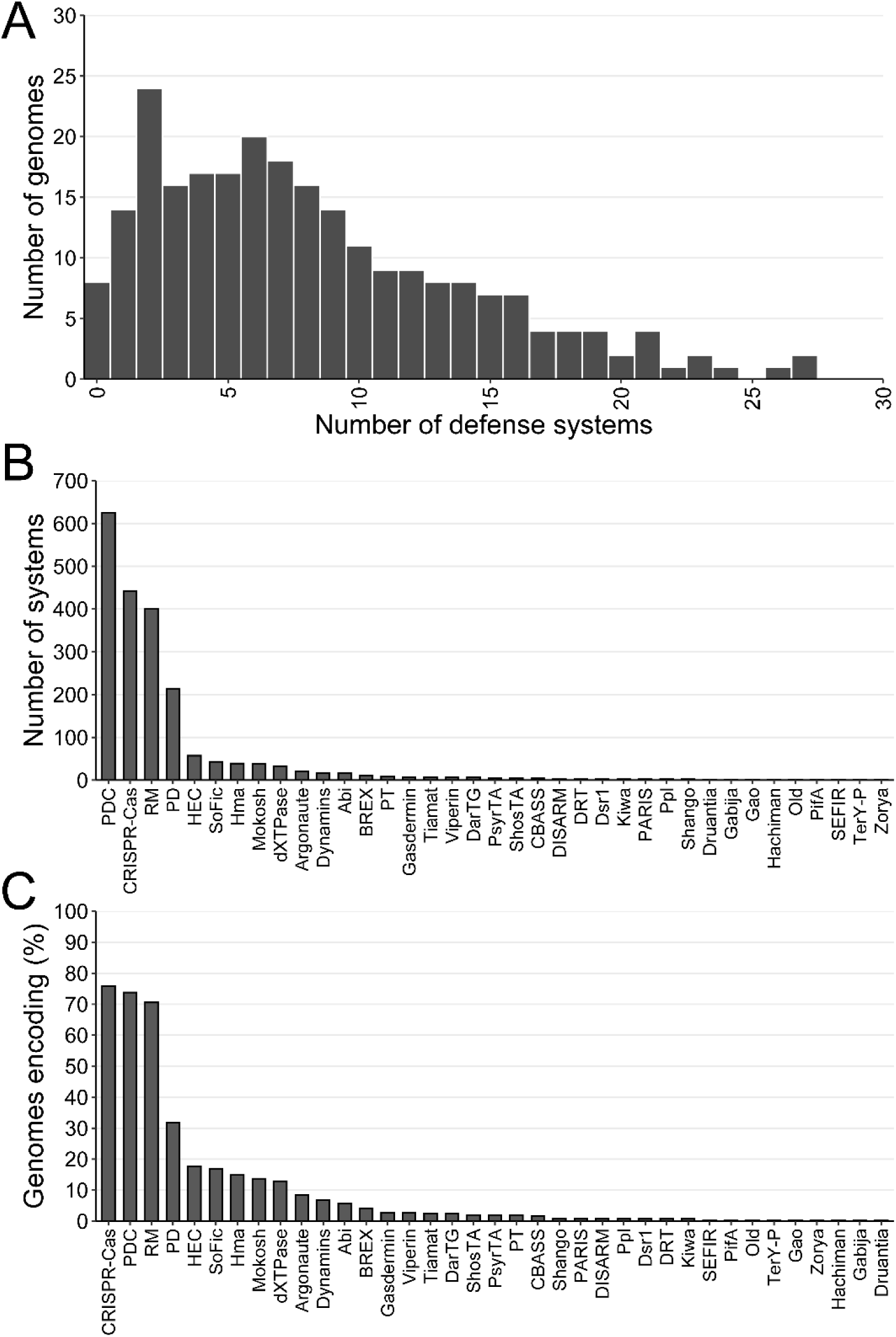
Families of defense systems in the prokaryotic genomes. (**A**) Total number of systems detected in pMAG/SAGs. (**B**) Distribution of the number of defense systems. (**C**) Frequency of systems.

To identify the MTases that catalyze the methylation of the detected motifs, MTase genes were systematically annotated (Figure S14; Data S3). Sequence similarity searches against known genes stored in REBASE^68^ identified 6.6 and 2.9 MTase and REase genes in prokaryotic genomes on average, respectively. Only 7 (2.8%) and 42 (16.9%) genomes lacked MTases and REases, respectively, suggesting that thermophilic microbes commonly possess RM systems, despite underestimation of defense system predictions owing to genome incompleteness. MTase genes were detected more than twice as often as REase genes across prokaryotic genomes, indicating that over half of the MTases were orphans. Across the M and RM genes in the prokaryotic genomes, m6A was the most abundant modification type, followed by m4C; m5C was almost absent, as observed in the HiFi read analysis (Figure 2F). Among the MTase types, Type II MTases were the most abundant, whereas a small number of the genes showed the highest sequence similarity to Type I and III MTases across the genomes. This trend was also consistent with the HiFi read analysis, which showed that Type I and III MTases were scarcely detected in the communities (Figure 2E). Overall, these analyses highlighted previously unknown MTases in hot spring biofilm communities, characterized by an overabundance of Type II MTases and a paucity of m5C modification types. In addition to the widespread RM systems supported by abundant REases and MTases in genome, the existence of numerous orphan MTases suggested unexplored roles for methylation systems beyond defense mechanisms.

A total of 291 (17.8%) MTase genes in prokaryotic genomes showed the highest sequence similarity to MTases, the specificity of which exactly matched the motif identified in our metaepigenomic analysis (Data S2 and S3). For example, a prokaryotic genome, pMAG_1st_NSBL_001, contained five MTase genes, three of which showed the highest sequence similarity to those that recognize **C**CG<u>G,</u> G**A**N<u>T</u>C, and G<u>T</u>**A**C, and this finding was perfectly congruent with the motifs detected in the pMAG. For example, pMAG_1st_NSBR_Circle_004 harbored 29 MTases, 13 of which were similar to those that recognize nine unique motifs (i.e., <u>T</u>CG**A**, G**A**N<u>T</u>C, G**A**<u>T</u>C, **C**TA<u>G</u>, <u>T</u>TA**A**, A<u>G</u>**C**T, C**A**<u>T</u>G, A**A**<u>T</u>T, and G**A**TA<u>T</u>C), which were congruently detected in the genome. In addition, one MTase gene was annotated as having GGCC (m4C) specificity, with the modification position yet to be determined; whereas, the metaepigenomic analysis identified <u>G</u>GC**C** in the same genome, suggesting that this motif is an actual specificity. Similarly, the pMAG_1st_SY_018 genome contained 38 MTases, eight of which showed the highest similarity to those that recognize the motifs exactly matching the predicted motifs on the same genome (i.e., G**A**CG<u>T</u>C, G**A**TA<u>T</u>C, G**A**N<u>T</u>C, C<u>T</u>GC**A**G, G**A**<u>T</u>C, C**A**<u>T</u>G, and G<u>T</u>CG**A**C). All the methylated motifs detected in the genome matched completely, suggesting that the MTases were active. In contrast to the congruent MTase-motif pairs, many MTases and detected methylated motifs were difficult to match. In 91 (36.7%) prokaryotic genomes, at least 1 MTase gene was found, however no methylated motifs were detected. We anticipate that either the MTase genes were inactive or the corresponding methylated motif was undetected owing to the low sensitivity of SMRT sequencing^26,73^.

In contrast to prokaryotes, MTase and REase genes were less frequently predicted, found in 131 (39.4%) and 18 (5.4%) viral genomes and in 182 (39.1%) and 86 (18.5%) extrachromosomal circular genomes, respectively. Compared with prokaryotes, M genes had a higher proportion of RM system genes in viral genomes (Figure S14; Data S3). Type I MTases were present in lower proportions in both viral and extrachromosomal circular genomes than in prokaryotes, likely because of the small genome size of mobilomes, which prefer not to maintain a larger Type I RM system (requiring at least one M and one S subunit gene) than Types II and III (one M gene). In addition, because REase would cause a rapid toxic effect on the host immediately after infection, viruses may tend to lack REase and REase-MTase hybrid genes in their genomes to ensure genome replication and assembly for propagation. These trends were almost concordant with those of a previous survey of genomes in the RefSeq database^22^. The number of MTase genes on vMAGs and eccMAGs showed no clear association with the presence of methylated motifs or with phylogenetic topology (Figures S10 and S11).

### Interactions between prokaryotes and mobilomes

We computationally predicted the hosts of viruses and extrachromosomal circular genomes to elucidate the potential interactions between prokaryotes and mobilomes within the community. Notably, genome-based prediction of phage–host interactions utilizes multidimensional signals, such as genome sequence similarity between hosts and mobilomes, and CRISPR spacer sequences, which record historical infections and thus do not always reflect current infections. Based on the genomic content, 1480 and 2960 possible relationships, accounting for 189 (56.9%) and 296 (63.7%) of vMAGs and eccMAGs, respectively, were predicted (Figures 5A and S15A). Among these interactions, 1161 (78.5%) and 2219 (75.0%), respectively, were predicted to involve pMAG/SAGs as the hosts, whereas the others were matched to GenBank genomes. Nineteen vMAGs were predicted to infect two different phyla, implying a broad host range, crossing phyla similar to those recently reported in hydrothermal mat microbial communities^78^. Similarly, 30 eccMAGs were predicted to belong to more than two phyla. Counting with duplicates, Aquificota was the host phylum most vMAGs possibly infect (56 vMAGs), followed by Deinococcota (30 vMAGs), Desulfobacterota (26 vMAGs), Acidobacteriota (24 vMAGs), and Chloroflexota (17 vMAGs). A few vMAGs were assigned to phyla that are currently not well characterized, such as HKB111 (1 vMAG), Hydrogenedentota (1 vMAG), Patescibacteriota (1 vMAG), Spirochaetota (1 vMAG), and WOR-3 (1 vMAG). Phages infecting Cyanobacteriota were rarely predicted, despite their high abundance (Figure 2B). This may have been due to the low in silico detection power for viral genomes from contig sequences or to other types of double-stranded DNA (dsDNA) phages (i.e., single-stranded DNA [ssDNA] and RNA phages), which are difficult to detect with the sequencing methods used in the present study. In case of eccMAGs, Aquificota was the most dominant host phylum (123 eccMAGs), followed by Deinococcota (49 eccMAGs), Chloroflexota (36 eccMAGs), Desulfobacterota (34 eccMAGs), Pseudomonadota (21 eccMAGs), and Acidobacteriota (9 eccMAGs). Furthermore, some eccMAGs were estimated to belong to a poorly characterized phylum, such as DUMJ01 (3 eccMAGs), DRYD01 (1 eccMAG), HKB111 (1 eccMAG), and WOR-3 (1 eccMAG).

**Figure 5.**
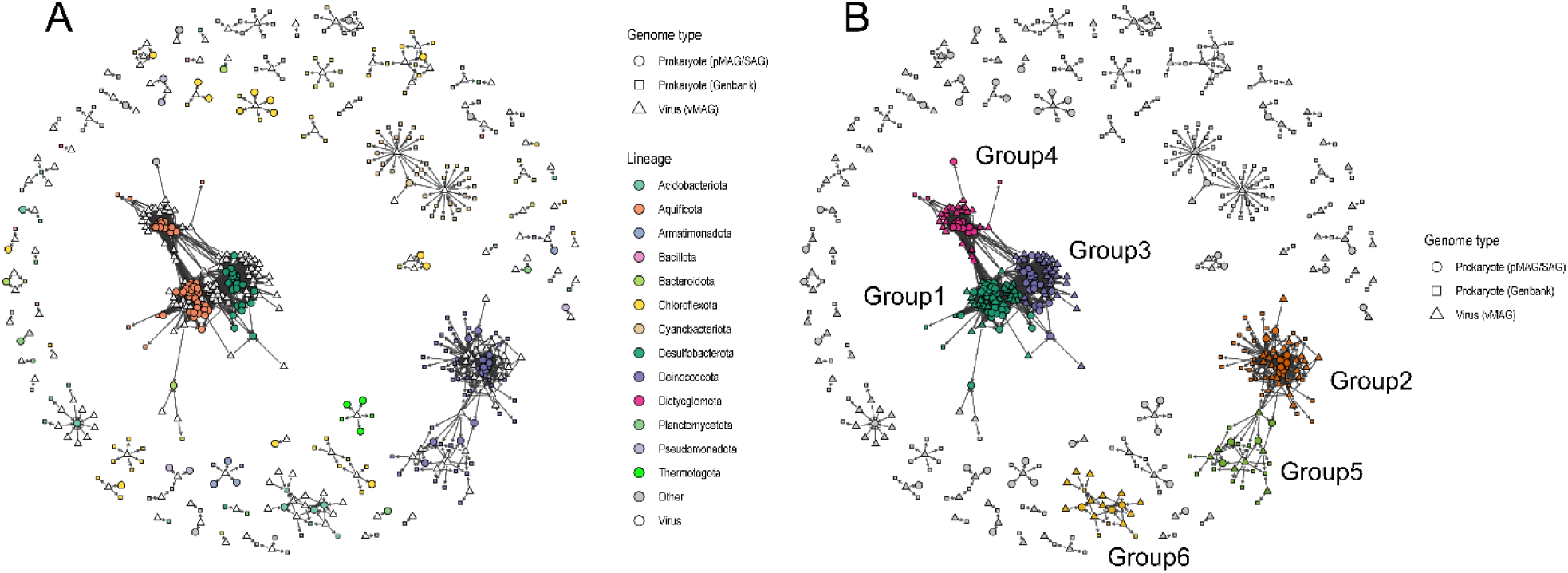
Network of virus–host interaction. (**A**) Phages and their prokaryotic hosts are shown as nodes. vMAGs, pMAG/SAGs, and prokaryotic genomes from the GenBank database are indicated by triangles, circles, and rectangles, respectively. Arrows represent infections of host prokaryotes by phages. Major prokaryotes are colored by taxonomic assignment at the phylum level. (**B**) Network colored by subnetworks, which were computationally determined. Nodes in the large subnetworks containing > 3 pMAG/SAGs are colored by group.

In the interaction network of vMAGs and their predicted hosts, complexity was associated with the host lineage (Figure 5A). The largest complex subnetwork comprised Aquificota, Desulfobacterota, and their phages, suggesting that many broad-host-range Aquificota/Desulfobacterota phages were present at the hot spring sites. The subnetwork associated with Deinococcota was also large, although its relative abundance was low across all the samples (Figure 2B). In sharp contrast, despite their high relative abundances, Chloroflexota and Acidobacteriota formed small, segmented subnetworks with their phages, suggesting that the host range of phages was narrow. Similarly, the network of extrachromosomal circular genomes showed several large subnetworks associated with the genomes of Aquificota, Desulfobacterota, and Deinococcota, whereas segmented subnetworks associated with Chloroflexota and Acidobacteriota were observed (Figure S15A).

Based on the potential interactive linkages between mobilomes and their hosts, we further investigated epigenomic similarities. Interaction subnetworks containing at least three pMAG/SAGs were examined, and it was found that the epigenomic patterns showed similarity within each subnetwork (Figures 4B and 5). In the case of viruses, for the largest prokaryotic subnetwork (Group 1), over half of the epigenomes shared several specific motifs (e.g., **C**CG<u>G</u>, T**C**<u>G</u>A, G**A**N<u>T</u>C, G<u>T</u>**A**C, and G**A**<u>T</u>C) across Aquificota and their phages. A large proportion of Group 2 showed similar trends, but different motif combinations, with **C**TCGA<u>G</u> and <u>T</u>CG**A** being comparatively shared among the genomes. The other motif set, **C**CG<u>G</u> and G**A**<u>T</u>C, was commonly shared across the genomes in Group 4. The motifs <u>T</u>TA**A** and <u>T</u>CG**A** were spread across the genomes in Group 5. In contrast, Group 6 comprised genomes that had low motif methylation ratios overall, although some encoded several MTase genes. In all cases, cluster analysis revealed mixed distributions of prokaryotes and phages, indicating that the phages and their hosts shared methylated motifs within each group. Similar trends were observed for extrachromosomal circular genomes (Figures S15B and S16). These were likely caused by infection, during which the mobilome genomes were modified by host-encoding MTases in the cell, as previously experimentally verified using pairs of cultured prokaryotes and isolated phages^79,80^.

Within each group, many methylated motifs were not evenly methylated but were clearly distinguished into several subgroups. For example, for viruses, Group 1 showed at least three large subgroups: the upper and middle subgroups showed more motifs with higher methylation ratios, whereas the lower subgroup was comparatively less methylated (Figure 6). Group 2 showed similar trends and was separated into two subgroups: the upper subgroup showed several conserved methylated motifs, whereas the lower subgroup lacked motifs with high methylation ratios. Group 3 was approximately divided into four subgroups. Group 4 comprised three subgroups with low and high numbers of methylated motifs in the upper and lower groups, respectively, except for the middle group, which consisted of only one prokaryotic genome and showed a different epigenomic pattern. Such epigenomic unevenness within each interaction network group also appeared in extrachromosomal circular genomes (Figure S16). This unevenness was not solely explained by prokaryotic phyla, sampling sites, or sequencing coverage. We anticipated that the unevenness reflected current infections in the communities, in contrast to the genome-based host prediction, which includes historical infections and mispredictions, considering that host prediction remains a challenging bioinformatics task^81^.

**Figure 6.**
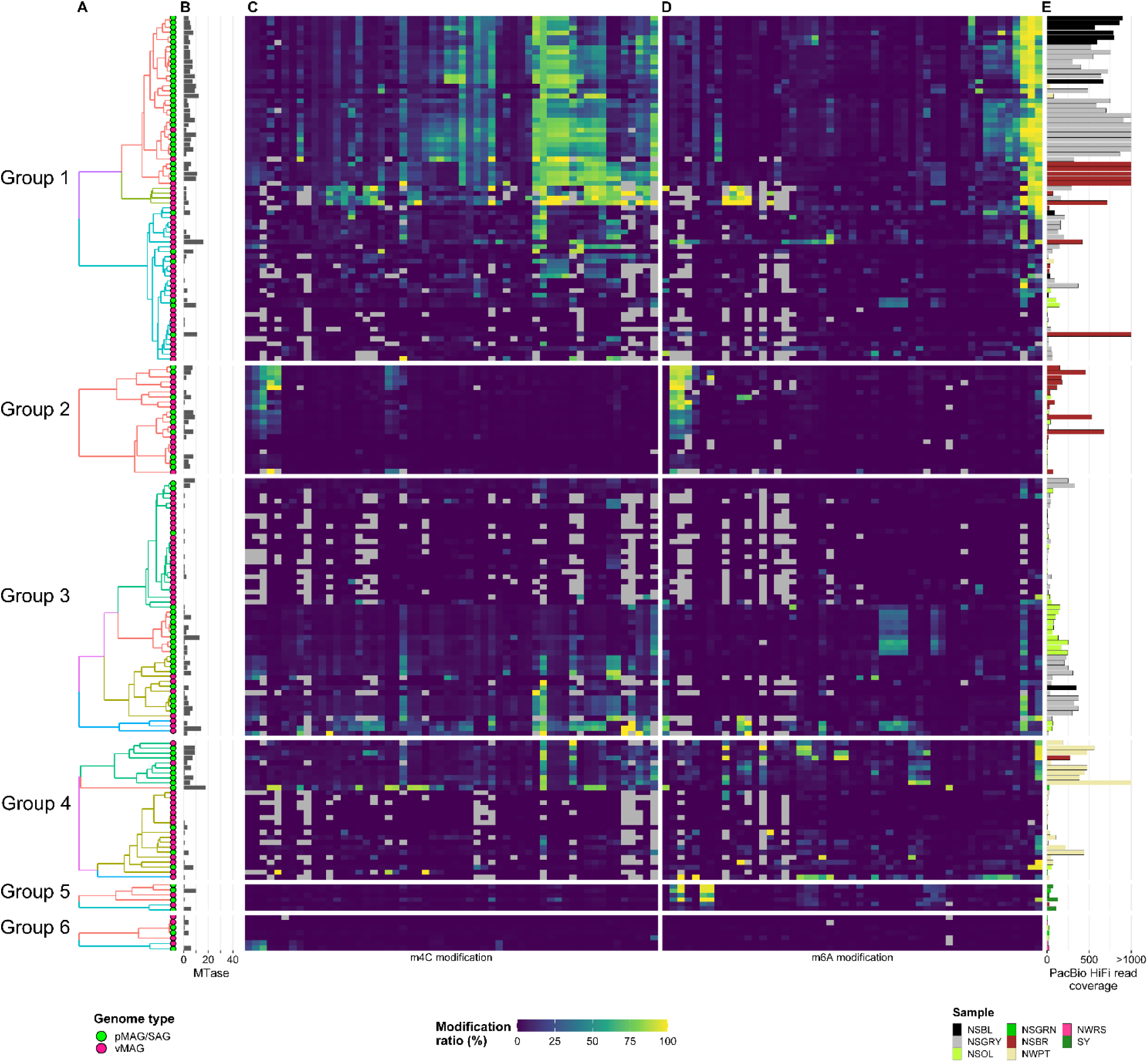
Methylomes of prokaryotic and viral genomes in each infection subnetwork. Hierarchical clustering of the modification ratio profiles within each subnetwork was performed independently using the Bray–Curtis dissimilarity index. The clusters in each group are colored in the dendrogram. Modification ratios of (**C**) m4C and (**D**) m6A motifs. The order of the motifs is the same as that shown in Figure 3.

## DISCUSSION

### Effect of DNA methylation in host–mobilome interactions

In this study, we first applied a single-cell genomic approach to metaepigenomic analysis and showed that diverse methylated motifs and defense systems were widespread across prokaryotes in hot spring biofilms. Almost half of the detected MTase genes in pMAG/SAGs were adjusted with a cognate REase gene or associated with known physiological systems such as BREX^76^ and DISARM^77^. This implied that some MTases were active in protecting against extracellular DNA invasion, whereas others were likely orphans. Because RM system genes in the communities shared similar compositions in terms of enzyme and modification types across the samples, the effects of environmental factors surrounding the hot springs (e.g., water chemical concentrations and temperature) may have been limited to the epigenome, in sharp contrast to the significant effects on the community composition (Figure 2). The relative abundance of RM system genes was higher than that in our previous analysis targeting marine water communities^34^, suggesting a higher infectious pressure in the biofilm than in planktonic habitats, possibly due to frequent contact with prokaryotic cells and viral particles, as supported by the high biomass density in the biofilm in the present study. This was also supported by the highly abundant non-methylation-mediated defense systems, such as the CRISPR-Cas system. Note that unprecedented defense systems associated with or independent of DNA modifications may have existed but were not detected in our analysis. Indeed, a number of novel defense systems have recently been proposed^82^ and may be discovered in the future. Therefore, further examination is required to identify comprehensive defensomes in these communities.

Methylated motifs in the hot spring prokaryotic genomes varied (Figure 3), whereas methylated ratios in the genomes and metagenomes were almost similar and concordant with those in the common *E. coli* DH5α (Figures S9). It has been anticipated that many prokaryotic lineages occasionally exchange RM systems within populations and that the methylated motifs associated with active RM systems have diversified to reinforce defense against extracellular DNA invasion over the course of the microbial arms race. Simply maintaining dozens of RM systems may help protect against any mobilome infection; however, this seems unrealistic because the more methylated bases in the genome, the greater the likelihood that they affect gene expression, leading to harmful physiological effects. Therefore, it does not follow a simple additive model. Thus, DNA methylation likely remains at low levels across diverse microbes, as observed in the present study. Supporting this, except for specific lineages such as *Helicobacter pylori*, in which over a dozen RM systems can be observed in a single genome^83^, the number of RM systems generally ranged from 0–6^22^, and the system abundances in hot spring prokaryotes revealed in the present study concordantly ranged from 0–9 (1.6 on average) (Data S1).

Alternatively, sets of a few of these systems with different combinations and diversified specificities would be an effective way to overcome viral infection, as phages need to be protected from as many host RM systems as possible to infect hosts effectively. Members with small numbers of methylated motifs in various combinations were observed in Chloroflexota, Bacteroidota, and Armatimonadota (Figure 3), which can be referred to as an ‘exchange’ strategy. The members exhibited small, isolated, infectious interaction subnetworks, suggesting that the diversified RM system increased resistance and limited the host range of phages (Figure 5). In contrast, those with very low motif diversity (‘retention’ strategy) were conserved in Aquificota, Deinococcota, Desulfobacterota, and Planctomycetota, which exhibit large, complex subnetworks. The fixed methylated motif profiles suggested a nearly identical set of active RM systems; hence, phages readily evaded uniform RM systems, resulting in their wide host range. It was also anticipated that these phyla are hot-spot organisms for genetic exchange at the hot spring sites. Indeed, some genera in Aquificota and Deinococcota have been reported to exhibit frequent population-wide recombination within their respective clades in hot spring environments^39^. Concordantly, the genomes of Aquificota, Deinococcota, and Desulfobacterota identified in this study showed very close phylogenetic relationships (short branches) within a few clades, possibly indicating genomic heterogeneity rather than speciation. In addition to defense systems, it is anticipated that some MTase genes regulate essential biological functions, resulting in high conservation within specific clades. This is similar to cell cycle-regulated MTases, which are highly conserved among members of Alphaproteobacteria^34,84^. Consequently, this sustained strategy is likely to be a driving force of rapid genomic evolution and epigenetic regulation of essential biological functions as a trade-off for a weak immune system. These two motif diversification strategies may be extended to other non-thermophiles; hence, it would be interesting to investigate a broader range of microbes.

Among viruses and extrachromosomal circular genomes, methylomes showed sporadic variation, and vMAGs and eccMAGs tended to lack MTase genes or have fewer than those observed for methylated motifs (Figures S10 and S11). Hence, it was suggested that DNA modification in mobilomes was mostly catalyzed by host-encoding MTases during invasion. Viral DNA modification mediated by host MTases has been examined in recent studies, and host and infectious phages share similar methylated motifs in their genomes, as determined by culture-based phage infection experiments using *Helicobacter pylori*^79^ and *Staphylococcus* isolates^80^. Consistent with these studies, we also found similar epigenomic patterns between mobilomes and their host prokaryotes (Figures 6 and S16). The shared epigenomic pattern suggested that mobilomes actually infected and resided in host prokaryotic cells (i.e., present infection) rather than a historical infection recorded in genomic regions such as spacers in the CRISPR array. This suggests the potential for further application: using epigenomic information, it would be possible to predict current host–phage infection and mobilome–transfection, rather than merely relying on genomic data that may be contaminated by remnants of past infections. It would be interesting to use epigenomic signals to predict the host of wide mobilomes.

RM system genes, including orphan MTase genes, are mobile and are often found on mobile elements, including phages and plasmids, likely resulting from gene transfer from the host chromosome^85^. Similar to the RM system, some phages and plasmids encode other defense systems, such as the CRISPR-Cas system, in their genomes^86,87^. Mobilome-encoding defense and DNA modification systems would affect the evolution and distribution of carrying mobilomes and regulate gene transfer between host genomes. In addition to genetic exchange, defense systems on mobilomes may help protect their hosts against superinfection by competing mobilomes, a phenomenon known as interviral competition^88^. Warfare within mobilomes in a community may play an important role in shaping gene flow dynamics across microbiomes and in the rise and fall of community populations.

### Potential alternative roles in DNA methylation across thermophilic communities

In addition to the defense system, methylation systems may play a role in gene regulation and DNA repair. During DNA replication in *E. coli*, DNA methylation functions as a marker of the original (parental) DNA strand and dictates mismatch repair on newly synthesized (daughter) unmethylated strands, a process known as methyl-directed mismatch repair^4,5^. High temperatures can compromise DNA stability; therefore, rapid and accurate DNA repair is crucial for survival in hot spring environments. In addition, because many phototrophic microbes are present in hot spring biofilms^72,89^, UV radiation may affect microbial DNA, similar to that in other aquatic environments. It is anticipated that DNA methylation may play a role in DNA repair, helping organisms adapt to thermal aquatic environments. However, further experimental and proteomic analyses are required to confirm the epigenetic regulation of the genes involved. In addition, we cannot exclude the possibility that methylation facilitates physicochemical DNA stability in high-temperature environments, as DNA methylation has been reported to increase the DNA melting temperature^90,91^.

Notably, hot spring biofilm microbes overall lacked m5C modifications in their genomes, as supported by MTase gene annotation (Figure 2F), epigenomic analysis (Data S2), and nucleoside analysis with LC-MS/MS (Figure S8). Consistent with our findings, m5C modification has been reported to be rarely observed in thermophilic bacterial isolates^92^. This trend is in sharp contrast to previously analyzed eukaryotes and many prokaryotes, including *E. coli* and marine metagenomes^34,93^, suggesting the universality of the m5C modification system through a wide range of environmental microbes, except for thermophiles. We anticipated that high temperatures would affect the thermostability of the m5C-type MTase or the m5C base, in which methyl residues are deaminated to thymine residues by heat, rendering them unusable in thermophiles.

### Challenges of experiments and bioinformatics in environmental metaepigenomics

Experimental validation is a crucial step in linking modified motifs identified by epigenomic analyses to modification systems identified by metagenomic analyses, and in determining each modification system and its specificity. For MTases, conventional *in-vivo* methodologies rely on heterogeneous protein expression in *E. coli* and methylation-sensitive restriction assays, which suffer from cytotoxicity, low throughput, and limited assay conditions; whereas, *in-vitro* enzyme assays based on protein purification require substantial effort and cost. An *in-vitro* one-pot assay using a reconstituted cell-free expression system that we previously developed enables rapid high-throughput screening, and is therefore an ideal approach for evaluating community-wide MTase genes^94^. However, no detectable MTase activities have been experimentally confirmed using either *in-vitro* or *in-vivo* approaches for multiple promising MTase genes and motifs identified from hot spring communities, indicating technical difficulties associated with thermophilic MTases (Methods S1; Note S4). Methodological improvements in the assay would bridge the gap between epigenomic phenomena and the molecular systems surrounding DNA modification, and thus be key to elucidating these systems in nature, including in extreme environments.

Long-read metagenomics is currently gaining increasing attention. Indeed, as in the present study, many studies have reported MAGs with higher connectivity and quality using metagenomic long reads across diverse environments, which are generally difficult to obtain using conventional short-read sequencing technologies alone. However, there remains considerable room for improvement in both experimental methodology and data analysis. In this study, we encountered widely used binning tools, all of which were designed for short reads, making them difficult to apply to our long HiFi read dataset. This was most likely due to differences in read length and contig connectivity driven by the comparatively low taxonomic diversity in extreme thermophilic environments and a rich set of sequencing reads, distinguishing it from typical metagenomic study settings. In addition, we encountered lineages with multiple short-repeat regions in their genomes, shaping significant heterogeneity within clonal populations and making it difficult to obtain representative assembled contigs using only long reads (Note S3; Figure S5). We further realized that multipartite genomes comprising multiple chromosomes in a single organism were difficult to bin into a single MAG using only long reads despite the reconstructed contigs being circular (Note S3). These issues may not be captured in conventional short-read-based metagenomic analyses, which generate shorter, unconnected contigs, highlighting the specific challenges of long-read metagenomics in low-biodiversity samples. Lineages whose genomes are difficult to reconstruct using only long reads, yet for which it is possible to obtain highly fragmented MAGs with short reads, may be widespread in nature and could be problematic in the era of long-read metagenomics.

Current SMRT sequencing technology supports a limited number of DNA modifications with sufficient reliability (i.e., m4C and m6A), although several DNA modifications occur in nature^2^. In addition to DNA modifications, RNA modifications have also received attention. Similar to dsDNA viruses and prokaryotes, DNA/RNA modifications are believed to play significant roles in ssDNA and RNA viruses. Nanopore sequencing technology is a potential approach for detecting both DNA/RNA modifications^95,96^, allowing broad modifications to be measured beyond limited methylations, as well as to examine the epigenomes of dsDNA and RNA viruses. However, because the sensitivity of DNA modification detection is currently much lower than that of PacBio platforms^97^, further improvements are needed. It would be interesting to investigate the epigenomes of these overlooked microbes as well as the community-level diversity in a wide range of environments using a metaepigenomic approach.

## CONCLUSION

The crucial biological roles of prokaryotic and viral DNA modifications have long been emphasized; however, knowledge regarding their diversity, evolutionary history, or ecological roles, including their effect on biological interactions with prokaryotes and mobilomes, is limited. To the best of our knowledge, this study is the first culture-independent meta-epigenomic analysis of a thermophilic microbial community. We produced a number of MAGs and SAGs, including high-quality and near-complete genomes, from diverse prokaryotes and viruses and successfully identified numerous previously unreported modified motifs and MTases in their genomes. The distinguished epigenomic distribution was attributed to a scarcity of m5C modifications across the hot spring biofilm samples, as supported by MS analysis, likely due to the low thermostability of the modification system, providing evidence of habitat-dependent preference in DNA modification. In addition, potential epigenomic associations between prokaryotes and mobilomes, some of which were mediated by DNA methylation, provided insights into the significant role of DNA modifications in microbial ecology. Despite these findings, we encountered several technical challenges in the accurate genome reconstruction, methylated motif identification, and enzymatic characterization of MTases. Moreover, our findings suggested the potential application of epigenomic information for valid host prediction, which is more accurate than using only genome sequence data which are affected by remnants of historical infections. Further investigations of various environmental communities would deepen current understanding of the relationships among molecular functions, ecological benefits, and evolutionary impacts of DNA modifications in diverse prokaryotes and mobilomes, as well as their interactions within communities. To achieve this, further methodological developments in sequencing technology, accurate assembly and binning tools, reliable detection of DNA/RNA modifications at single-base resolution, motif prediction, and experimental validation of the modification system are warranted.

## DATA AVAILABILITY

The metagenomic HiFi sequencing data were deposited in the DDBJ Sequence Read Archive (Table S3). The prokaryotic and viral genomes were deposited in DDBJ/ENA/GenBank (Data S1). The data are registered under BioProject ID PRJDB42039 [https://ddbj.nig.ac.jp/resource/bioproject/PRJDB42039]. Extrachromosomal circular genomes were deposited in Zenodo [https://doi.org/10.5281/zenodo.20755168]. The scripts used in the study are available online under a CC-BY-NC license at https://github.com/hiraokas/Metaepigenome.

## ACKNOWLEDGEMENTS

We are grateful to Takahito Momose (the owner of Nakabusa hot spring) for allowing us to stay and collect samples from the hot spring sites. We also thank Katsumi Matsuura for the joint collection of microbial biofilms. Finally, we would like to take this opportunity to thank Hiroshi Shimizu, Hideyuki Tamaki, and Nobuhiko Nomura for their thoughtful advice and encouragement. Bioinformatics analysis was performed using the supercomputing system at the National Institute of Genetics (NIG), Research Organization of Information and Systems (ROIS), and the Earth Simulator systems at JAMSTEC.

## Author contributions

SH conceived and designed the study, performed the sampling, molecular experiments, bioinformatics analyses, and wrote the manuscript. SS performed the mass spectrometry analysis. KU and TS designed and performed the molecular experiments, protein purification, and *in vivo*/*in vitro* MTase assays using heterogeneous expression. YZ performed mass spectrometry analysis and the *in vitro* MTase assay using the one-pot approach. YT performed sampling. SK and AN performed sampling and chemical measurements. All authors read and approved the final manuscript.

## FUNDING

The Japan Society for the Promotion of Science (JP19K16234, JP19K21203, JP20J13082, JP20K15444, JP20K21455, JP22K05398, and JP25K18446); The Japan Science and Technology Agency, ACT-X (JPMJAX22BK); The Institute for Fermentation, Osaka (IFO).

## Conflict of interest statement

None declared.

## Notes

### Competing Interest Statement

The authors have declared no competing interest.

