## SUPPLEMENTARY INFORMATION for "Epigenomes in thermophilic microbial communities and their impact on the interaction across prokaryotes and mobilomes"

This PDF file includes:

- Supplementary Methods S1
- Supplementary Notes S1 to S4
- Supplementary References
- Supplementary Figures S1 to S16.
- Supplementary Tables S1 to S5.
- Supplementary Data S1 to S4.

### Supplementary Methods

**Methods S1. Experimental verification of MTase activities**

To verify MTase specificity, we used an *in-vitro* one-pot characterization approach[1], an *in-vivo* REase assay using *E. coli* heterogeneous expression, and an *in-vivo* REase assay using purified MTase. MTase genes were selected for artificial gene synthesis with codon optimization by Eurofins Genomics (Ebersberg, Germany).

In the one-pot approach, a series of overlap–extension polymerase chain reactions (PCRs) (Platinum SuperFi PCR Master Mix, Invitrogen, Waltham, Massachusetts, USA) were designed and used to prepare linear template DNA for the putative MTases. A universal DNA fragment (namely, Fragment 1) containing 5′ UTR (untranslated region, including T7 promoter and ribosome binding site) was PCR amplified (30 cycles) from a T7 expression vector pET-3a with a proper primer set (P1F and P1R). Another DNA fragment (Fragment 2), containing the CDS of the target protein, was amplified using the corresponding artificial gene as a template and an additional set of primers (P2F and P2R). The 5′ end of the P2R primer included 14 arbitrary nucleotides downstream of the stop codon, as recommended for linear DNA templates used in cell-free protein expression. Fragments 1 and 2 were confirmed and recovered using E-Gel CloneWell II Gel (Invitrogen). Since the 5′ ends of primers P1R and P2F were designed to overlap with each other, there was an overlapping region between Fragments 1 and 2. Equal amounts of these two fragments, with a total mass of approximately 10–20 ng, were mixed and subjected to short PCR (15 cycles) without additional primers. A 0.1-µL aliquot of the PCR solution was directly added to a fresh 50-µL PCR reaction containing forward primer P1F and reverse primer P2R. The PCR amplicons were confirmed by agarose gel electrophoresis. The PCR product was column purified using a QIAquick PCR Purification Kit (Qiagen) and quantified using a NanoDrop One (Thermo Fisher Scientific).

The *in-vitro* cell-free protein synthesis (CFPS) reaction was conducted in a standard microtube (FastGene 0.2-mL PCR tube) (NIPPON Genetics, Tokyo, Japan). The template DNA of the putative MTase (3 ng/µL per 1 kb DNA) was added to a reconstituted CFPS PUREfrex 2.0 (GeneFrontier, Chiba, Japan) containing 10 µL of Solution I, 1 µL of Solution II, 2 µL of Solution III, and nuclease-free $H_2O$ (NIPPON GENE, Tokyo, Japan) in a total volume of 20 µL. For putative MTases predicted by DIpro to form disulfide bonds[2], PUREfrex 2.1 (i.e., an altered version of PUREfrex 2.0 optimized for disulfide bond formation) was tested in combination with glutathione disulfide (GSSG) and the disulfide bond isomerase DsbC (GeneFrontier). For PUREfrex 2.1, the 20-µL reaction solution was composed of 8 µL of Solution I (a different composition without cysteine and reducing agents), 1 µL of cysteine, 1 µL of glutathione, 0.33 µL of GSSG, 0.25 µL of DsbC, 1 µL of Solution II, 2 µL of Solution III, template DNA (3 ng/µL per 1 kb), and nuclease-free $H_2O$. All CFPS reactions were performed at 37 °C for 4 h on a PCR thermocycler (Veriti, Applied Biosystems, Carlsbad, California, USA). Each CFPS solution was subjected to protein electrophoresis using an Agilent 2100 Bioanalyzer with an Agilent Protein 80 Kit to confirm the protein expression levels. The resulting CFPS solutions were stored at −80 °C until use.

MTase activity was detected using an MTase-Glo assay (Promega, Madison, Wisconsin, USA). Briefly, each CFPS reaction solution was dialyzed using a Micro Tube-O-DIALYZER, 15 K MWCO (G-Biosciences, St. Louis, Missouri, USA) against 0.1 M Tris-HCl buffer (pH 8.0) at 4 °C. A 1-µL aliquot of the dialyzed MTase sample was mixed with 10 µM of a hairpin DNA oligo substrate (synthesized by FASMAC) and 50 µM SAM (from the MTase-Glo assay kit) in 1× reaction buffer (20 mM Tris-HCl, pH 8.0, 50 mM NaCl, 1 mM EDTA, 3 mM $MgCl_2$, 0.1 mg/mL BSA, and 1 mM dithiothreitol) to a total volume of 4 µL. The DNA methylation reaction was carried out at room temperature (23 °C) for 10–60 min. Then, 1 µL of 5× MTase-Glo Reagent (from the kit) was added to the above reaction mixture, followed by an additional 30 min incubation at room temperature, which converted the methylation reaction by-product, S-adenosylhomocysteine, to ADP. Finally, the ADP was converted to ATP by adding 5 µL of MTase-Glo Detection Solution (from the kit) to the mixture, and ATP was detected using luciferase. The chemiluminescence reaction was performed at room temperature in a 384-well microplate (low-volume, white, non-binding surface; Corning, Tewksbury, Massachusetts, USA) for 30 min. An endpoint luminescence measurement was performed using a BioTek Synergy H1M2 plate reader (Agilent Technologies).

For the *in-vivo* assay, the MTase genes were cloned into the pCold III expression vector (Takara Bio) using an In-Fusion HD Cloning Kit (Takara Bio). If an appropriate sequence was absent from the plasmid vector, additional specific sequences were inserted downstream of the termination codon for the methylation assay. The constructs were transformed into *E. coli* HST04 *dam*$^-$/*dcm*$^-$ (Takara Bio), which lacks the Dam and Dcm MTase genes. Soluble protein concentrations were measured by SDS-PAGE. *E. coli* strains were cultured in LB broth supplemented with ampicillin. MTase expression was induced according to the supplier's protocol for the expression vector. Plasmid DNA was isolated using a NucleoSpin Plasmid EasyPure (Takara Bio). The methylation status was assayed simultaneously with linearization by digestion with the appropriate REases, and specific REases were employed to linearize the plasmid DNA. The REases were purchased from NEB and Nippon Genetics. All digestion reactions were performed at 37 °C for 1 h. DNA fragments were separated using agarose gel electrophoresis. We further verified the specificity of MTase via SMRT sequencing. The chromosomal DNA of *E. coli* HST04 *dam*$^-$/*dcm*$^-$ strains, in which the target MTase was transformed, was extracted using a DNeasy UltraClean Microbial Kit (QIAGEN), according to the supplier's protocol, after induction of gene expression. Multiplex SMRT sequencing was performed on a PacBio Revio system (Pacific Biosciences of California) according to the manufacturer's standard protocols. Methylated motifs were predicted using SMRT Link v25.3 against the *E. coli* K-12 MG1655 reference genome (RefSeq NC_000913.2).

For the *in-vitro* assay of the selected purified MTase, the N-terminal 6×His-tag fusion MTase mutant was constructed using PCR and cloned into the pCold II expression vector. *E. coli* HST04 *dam*$^-$/*dcm*$^-$ cells, transformed with the constructs, were grown at 37 °C for 16 h in 500 mL of medium A (LB medium containing 100 μg/mL ampicillin) via shaking. The culture was inoculated into 1 L of medium A in a 3-L flask (5 flasks; total 5 L), incubated at 37 °C for 2–3 h with constant shaking, and allowed to grow until the optical density at 600 nm reached 0.6. Then, MTase expression was induced with 0.1 mM isopropyl β-D-1-thiogalactopyranoside (IPTG) at 15 °C, and the cultures were subsequently incubated for 16 h. *E. coli* cells were harvested and suspended in Buffer A (20 mM HEPES-Na [pH 7.5], 150 mM NaCl, 5% glycerol, 1 mM DTT, and 50 mM imidazole). The suspension was lysed at 800 bar using a high-pressure homogenizer LAB2000 (SMT Co., Ltd.) for three passes. The cell lysate was centrifuged at 15,000 × g for 20 min at 4 °C, then passed through a 0.45-μm GD/X syringe filter (Cytiva, Marlborough, Massachusetts, USA). The supernatant was subjected to three-column chromatography using the ÄKTA start and ÄKTA go chromatography systems (Cytiva). The sample was loaded onto a 5-mL HisTrap HP column (Cytiva) at a flow rate of 2 mL/min. The column was then washed with Buffer A. The His-tagged protein was eluted with Buffer B (20 mM HEPES-Na [pH 7.5], 150 mM NaCl, 5% glycerol, 1 mM DTT, and 300 mM imidazole). The enzyme was further purified by ion-exchange (5-mL HiTrapQ column [Cytiva]) and size exclusion chromatography (HiLoad 16/600 Superdex 200 pg column [Cytiva]) using Buffer C (20 mM HEPES-Na [pH 7.5], 150 mM NaCl, and 1 mM DTT). The protein was collected as a single peak and concentrated to 0.878 mg/mL (approximately 8.3 μM monomer). It was then aliquoted, flash-frozen in liquid nitrogen, and stored at −80 °C until further use.

The purified MTase was used for enzymatic methylation. The substrate (i.e., unmethylated DNA) was amplified by PCR using the pCold II vector, which contained the same MTase gene as the template. Methylation reactions were carried out in a reaction buffer (20 mM HEPES-Na, pH 7.5, 100 mM NaCl, and 100 μg/mL BSA) containing 10 nM substrate DNA and 2.1 μM purified MTase at 70 °C for 1 h or 3 h. The reaction temperature was determined based on the sampling site temperature and the Tm value measured by the protein thermal shift assay using a StepOnePlus Real-Time PCR System (Applied Biosystems) with Protein Thermal Shift Dye (Applied Biosystems) and Protein Thermal Shift software (v1.4) (Applied Biosystems). The reactions were initiated with 160 μM SAM (NEB) in solution and terminated by adding NTI buffer containing guanidinium thiocyanate

(Takara Bio). After the methylation reaction and subsequent DNA purification, the methylation status was assessed via agarose gel electrophoresis and capillary electrophoresis using a 5300 Fragment Analyzer System (Agilent Technologies, Santa Clara, California, USA) and a HS NGS Fragment Kit (Agilent Technologies) after REase digestion at 37 °C for 1 h.

### Supplementary Notes

##### Note S1. Geochemical characteristics of the hot springs

Geochemical properties differed among the three hot springs (Table S2). Within the samples from the Nakabusa stream site, the concentrations of the measured items were similar; however, there were slight differences between the two streams, whereby $Cl^-$, $Na^+$, and $K^+$ concentrations were slightly lower, whereas the $Mg^{2+}$ concentration was higher, in one stream line (N-1–N-2) than in the other line (N-3–N-4) (Figure S2A). A notable exception was $H_2S$, which showed comparatively high concentrations upstream and significantly lower concentrations downstream of the current (below the detection limit at N-2 and N-5). This may have been associated with sulfide oxidation by dominant microbes (e.g., *Sulfurihydrogenibium* and *Chloroflexus*)[3,4] and/or abiotic sulfide oxidation driven by oxygen produced by oxygenic phototrophic Cyanobacteriota in biofilms, which has been previously investigated[5]. These processes may have also resulted in the relatively high sulfate concentration at N-5. The wall site at the Nakabusa hot spring showed similar chemical components, but with slightly higher $Cl^-$ and $Na^+$ concentrations and lower $SO_4^{2-}$ concentrations. Overall, the concentrations in Nakabusa hot spring were consistent with previous observations[5]. The source water of Shiriyaki hot spring exhibited moderate $SO_4^{2-}$ concentrations and low $Cl^-$ and $Na^+$ concentrations. Dewanoyu was a sulfidic hot spring characterized by high concentrations of salts and dissolved minerals, including $Cl^-$, $Na^+$, $K^+$, and $Mg^{2+}$, as well as a slightly elevated Mn concentration.

##### Note S2. Prokaryotic community structure of hot spring biofilms

Bacteria were dominant across the samples, whereas almost 0.1–2.3% of the HiFi reads were assigned to Archaea, except for NSBL and NSGRY, which were collected at the higher-temperature sites in this study and reached 23.6–26.4% of the HiFi reads (Figure 1A). The most dominant phyla in NSBL and NSGRY were Aquificota, followed by Thermoproteota and Thermodesulfobacteriota (Figure 1B). The NWPT community was dominated by Aquificota, Thermodesulfobacteriota, and Deinococcota. These community structures were concordant with previous reports based on 16S rRNA amplicon sequencing[6]. The most abundant phylum in NSBR was Aquificota, followed by Deinococcota, Bacteroidota, Armatimonadota, and Acidobacteriota. In contrast, Cyanobacteriota was the most abundant member in NSGRN, NSGF, and SY, all of which were green colored, likely because of the high chlorophyll *a* levels in the lineage. The Chloroflexia class, belonging to the phylum Chloroflexiota, was dominant in the NWRS, where the *Roseiflexus* genus was abundant (Figure 1C), and the biofilm was rose colored, resulting from the richness of the gamma-carotene derivatives utilized for photosynthetic pigments[7]. Chloroflexiota was also abundantly detected in NSOL and NSGRN, consistent with a previous report[8]. The DW sample exhibited a distinct community structure with abundant Pseudomonadota, Deferribacterota, and Elusi microbiota. In the samples from Nakabusa hot spring, the community composition was overall concordant with that reported in previous studies[5,6,8–16] (Table S1).

**Note S3. Genome reconstruction and taxonomic assignment**

The HiFi reads were assembled into 4899–32677 contigs from each sample using hifiasm-meta, which was optimized for metagenomic HiFi reads[17] in the first step of our assembly strategy (see Materials and Methods) (Figure S2; Table S3). The total length of the assembled contigs ranged from 209–1290 Mb, the N50 values were 19–65 kb, and the length of the longest contigs reached 186–560 kb. A total of 238 assembled graphs (ranging from 7–53 per sample) that showed complexity above our threshold were then subjected to a second assembly using hifiasm, an assembler for a single organism[18]. Gathering the contigs from the first and second assemblies, the contigs were binned into a total of 373, 332, and 465 for the prokaryotic (pMAGs), viral (vMAGs), and extrachromosomal circular genomes (eccMAGs), respectively (Data S1).

However, we found a lack of *Chloroflexus* genus genomes in the NSOL samples. The genus *Chloroflexus*, which belongs to the phylum Chloroflexota, is known as the most dominant bacterial member in the NSOL site[5,8,15] and was indeed highly abundant in NSOL, as estimated from the HiFi reads (Figure 1B). The absence of MAGs assigned to a specific genus was most likely not due to insufficient sequencing data, read quality, or bioinformatic methodology. One of the most dominant species in this field was *Chloroflexus aggregans*, a metabolically versatile, thermophilic, and anoxygenic phototrophic member of the phylum Chloroflexota[19]. We found that the type strain of this species (*Chloroflexus aggregans* DSM 9485 = MD-66 = JCM 39377) contained several SSRs on its complete genome (RefSeq accession ID: CP001337.1): at least 67 regions with > 10 times repeat composed with > 8 bp sequence units (Table S4). HiFi reads from NSOL mapped to SSR regions exhibited high heterogeneity, suggesting that repeat frequency varied at the cell level within the population (Figure S5). This may have been caused by the slipped-strand mispairings during DNA replication[20]. Owing to the significant heterogeneity, it was difficult to uniquely determine a single representative genome for this strain. In practice, highly complex bubble structures in the assembled graphs in such SSR regions are split during metagenome assembly, even when long reads span the entire region. Therefore, no Chrolofrexiota pMAGs were retrieved using the proposed binning strategy.

To obtain the missing genomes, we conducted single-cell genomics to recover SAGs using the NSOL sample. Using a droplet-based single-cell isolation approach and short-read sequencing following whole-genome amplification, we obtained 114 high- or moderate-quality SAGs. After de-replication of the total pMAGs and pSAGs, 248 strain-level representative genomes, including two *Chloroflexus* pSAGs (see below), were identified as pMAG/SAGs and used for further analysis.

The completeness of pMAG/SAGs reached 81.5% on average (Figure 6A). Notably, 80 genomes had circular structures with > 90% completeness, suggesting complete prokaryotic chromosomes. In contrast, five circular pMAGs showed comparatively low completeness (ranging from 81.8–89.6%). Among them, one genome, pMAG_1st_NSGF_Circle_002, was assigned to Chloracidobacterium, the members of which members harbor multipartite genomes comprising two circular chromosomes[21], implying that the corresponding smaller circular genome was missing from the MAG. In addition, because the tool used for genome quality estimation, CheckM2, is a machine learning-based method that relies on a reference genome dataset[22], it was anticipated that the completeness of thermophilic microbial genomes would be underestimated, given the paucity of genome resources relative to more general species. The estimated contamination levels were low overall (0.9% on average). The pMAG/SAGs spanned 38 phyla, including 3 groups that are not yet well characterized: DRYD01, DUMJ01, and WOR-3. The most abundant phyla were Aquificota (45 pMAGs and 1 pSAGs), followed by Desulfobacterota (17 pMAGs and 13 pSAGs), Chloroflexota (27 pMAGs and 2 pSAGs),

Patescibacteriota (also referred to as Candidate Phyla Radiation or CPR[23]) (19 pMAGs and 3 pSAGs), and Deinococcota (9 pMAGs and 5 pSAGs). As expected, the two Chloroflexota pSAGs were assigned to *Chloroflexus aggregans* at the species level. The pMAGs assigned to Archaea were composed of six phyla: Thermoproteota (14 pMAGs), Halobacteriota (1 pMAG), Korarchaeota (1 pMAG), and the DPANN superphylum (Aenigmatarchaeota [1 pMAG], Iainarchaeota [pMAG], and Micrarchaeota [1 pMAG]). The Halobacteriota genome, a halophilic archaeon, was consistently identified at the Dewanoyu site, which features a highly saline hot spring (Table S2).

The average completeness of vMAGs was 76.1% (Figure 7B). Among the vMAGs, 80 (24.1%) were retrieved from NSGRN, followed by NSGRY (49; 14.8%), and NSOL (47; 14.2%). In contrast, only five (1.5%), six (1.8%), and nine (2.7%) were retrieved from NSBL, DW, and NSGF, respectively. This difference may have reflected diversity and variation in the abundance of dsDNA viruses. CheckV[24] reported that 6 and 93 vMAGs met the complete and high-quality criteria, respectively. The length of the vMAGs ranged from 10–409 kb, with an average of 50 kb. Seventy vMAGs were identified as proviruses, likely benefiting from long, accurate metagenomic reads that produced long contigs. In total, 156 of 332 (47%) vMAGs were taxonomically assigned at least to the class level. Among them, Caudoviricetes were dominant, except for three Hukuchivirus and three Matsushitaviridae vMAGs.

Among the eccMAGs, 129 (27.7%) were retrieved from NSBR, followed by NSGRY (83; 17.8%) and NSGRN (69; 14.8%). In contrast, only 9 (1.9%), 16 (3.4%), and 17 (3.7%) were retrieved from NSGF, NWRS, and NSBL, respectively. This unevenness likely reflected differences in sequencing read sizes, given that we focused only on circularized genomes (Table S3), and further sequencing efforts are needed to obtain a comprehensive view of extrachromosomal circular genomes in the communities. The length of the eccMAGs ranged from 3–406 kb, with an average of 50 kb. In total, 123 (26.5%) were classified as plasmids.

HiFi read coverage using the same samples as the genome-derived were highly divergent, ranging from 3.0–5554.9×, 1.4–8524.4×, and 0.0–42871.7× per pMAG/SAGs, vMAGs, and eccMAGs, respectively. Among the pMAGs/SAGs, 161 of 248 genomes showed > 50× coverage of the double strand, which is likely to be sufficient to detect m6A and m4C modifications according to the official white paper (https://www.pacb.com/wp-content/uploads/2015/09/WP_Detecting_DNA_Base_Modifications_Using_SMRT_Sequencing.pdf), despite the fact that the values were estimated using outdated RSII platforms with the premodern Continuous Long Read (CRL) method.

**Note S4. Exploration of novel MTases and experimental verification of their specificities**

Among the detected MTase genes from prokaryotic genomes, 132 (74%) showed inconsistencies between the recognition motifs of their closest relatives and the methylated motifs identified in the metaepigenomic analysis (Data S2 and S3). This result supports the empirical rule that the homology-based estimation of MTase specificity was insufficient, as in our previous meta-epigenomic investigations of freshwater[25] and seawater[26] microbiomes. To determine the catalytic specificity of these MTases, we selected potential pairs of putative MTases and methylated motifs as follows: 1) the MTase and methylated motifs were present in the same prokaryotic genome, and a novel correspondence was predicted; 2) the modification type of the MTase and the corresponding methylated motif were concordant; and 3) the complete sequence of the MTase gene was retrieved. Using this criterion, 11 MTase systems (including one Type I MTase composed of M and S subunit genes) were selected and subjected to biochemical experiments (Table S5).

First, the methylation specificities of selected MTases were experimentally verified using an *in-vitro* one-pot approach[1]. Among the MTases, three were assayed at 60 °C, in addition to room temperature (23 °C), to match the temperatures at the sampling sites. Although the positive control, MTase, showed clear positive results, no MTase activity was detected in the selected proteins from the hot spring biofilm samples, which may have been because the expression or assay conditions were far from optimal.

Next, we conducted conventional *in-vivo* experiments using three selected MTases via heterologous expression in *E. coli*. Briefly, plasmids containing a single artificially synthesized MTase gene were constructed and transformed into *E. coli* cells. The methylation status of the isolated plasmid DNA was then assessed by REase digestion following heterologous expression. However, this trial also failed, making it difficult to determine its specificity, except for one (pMAG_1st_NSBR_Circle_003_1829) which showed faint modification signals. Through resequencing analysis of the *E. coli* genome in which MTase was induced using SMRT sequencing, we failed to determine its *de novo* specificity, which may have been because of low methylation activity. Further, *in-vitro* enzymatic analysis of the purified MTase protein was conducted; however, no clear inhibition of REase digestion by DNA methylation was observed.

### Supplementary Figures

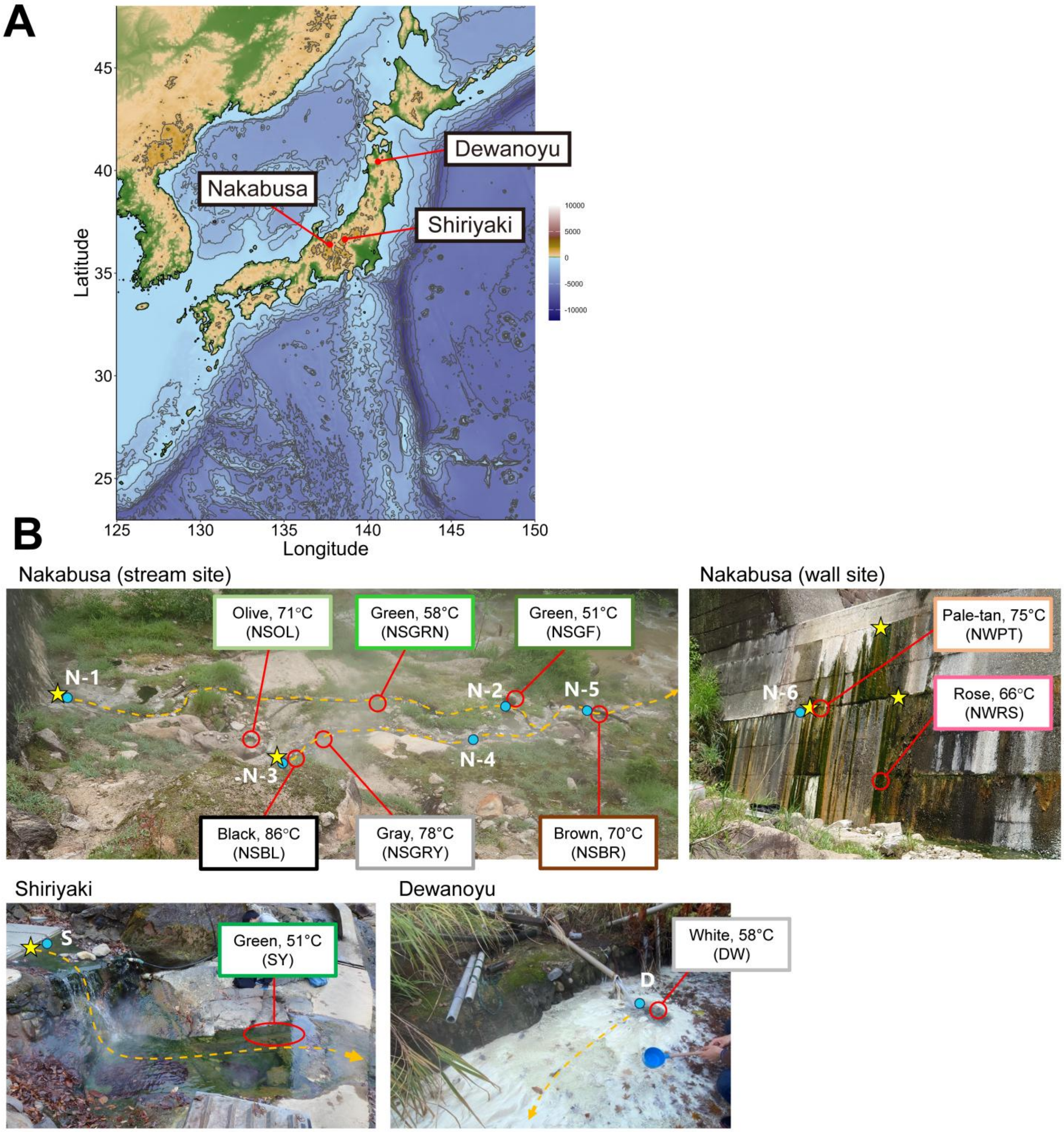


**Figure S1. Sampling site locations and photographs.**

(**A**) Map of the hot springs. (**B**) Photographs of the sampling sites. Sampling spots for microbial analysis and water chemical measurements are represented by red open and blue circles, respectively. Yellow stars indicate major sources of the hot spring water. Orange break lines represent major water currents.

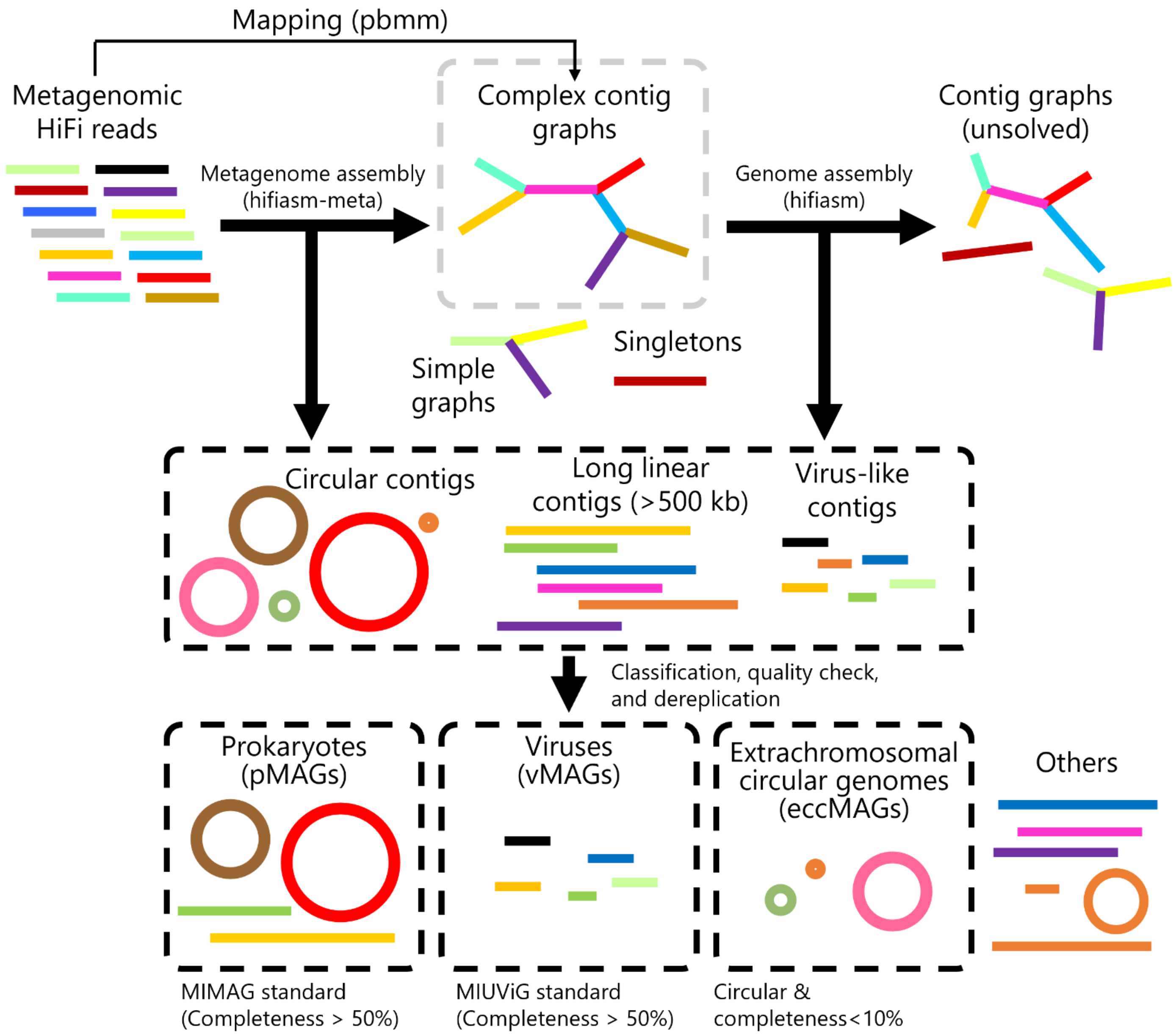


**Figure S2. Schematic view of the metagenome assembly and MAG construction workflow.**

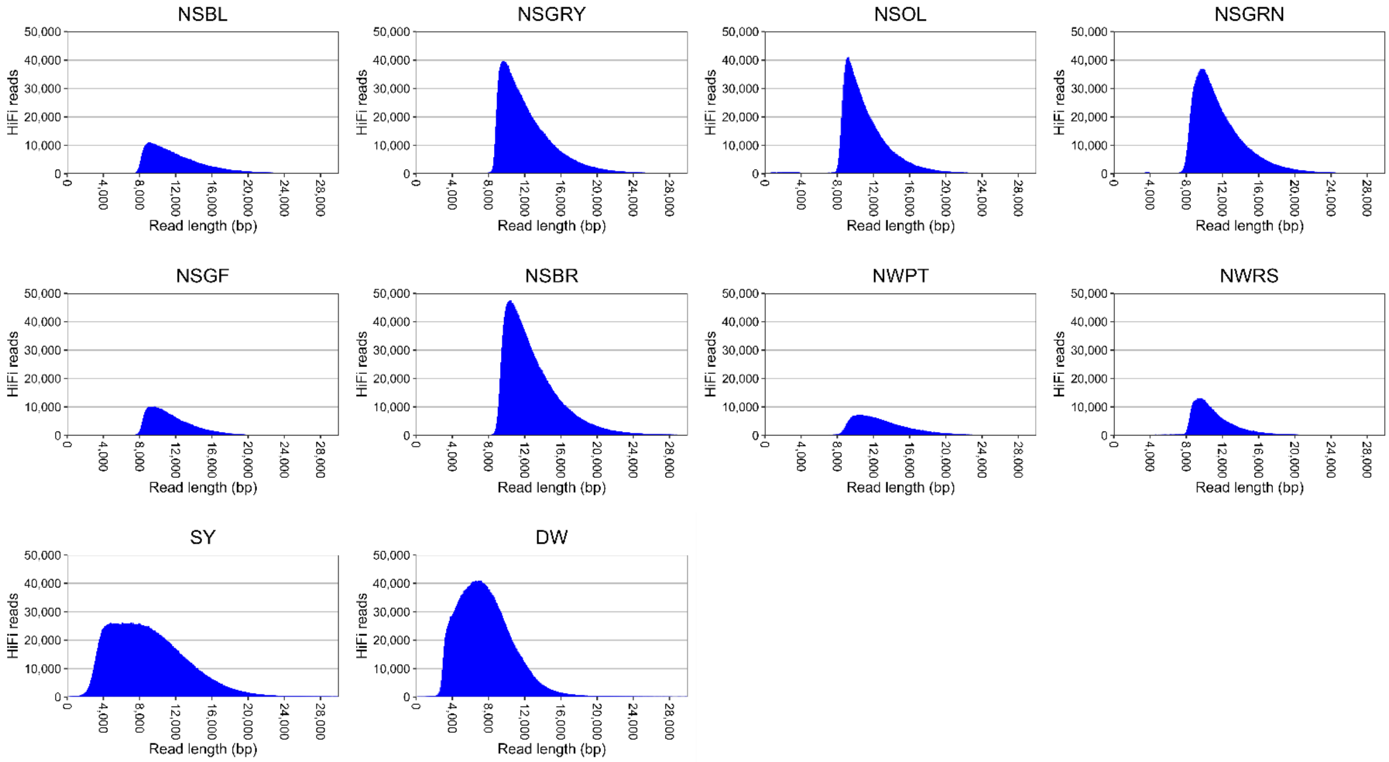


**Figure S3. Frequency distribution of the length of metagenomic HiFi reads.**

Read lengths were binned in 100-bp increments.

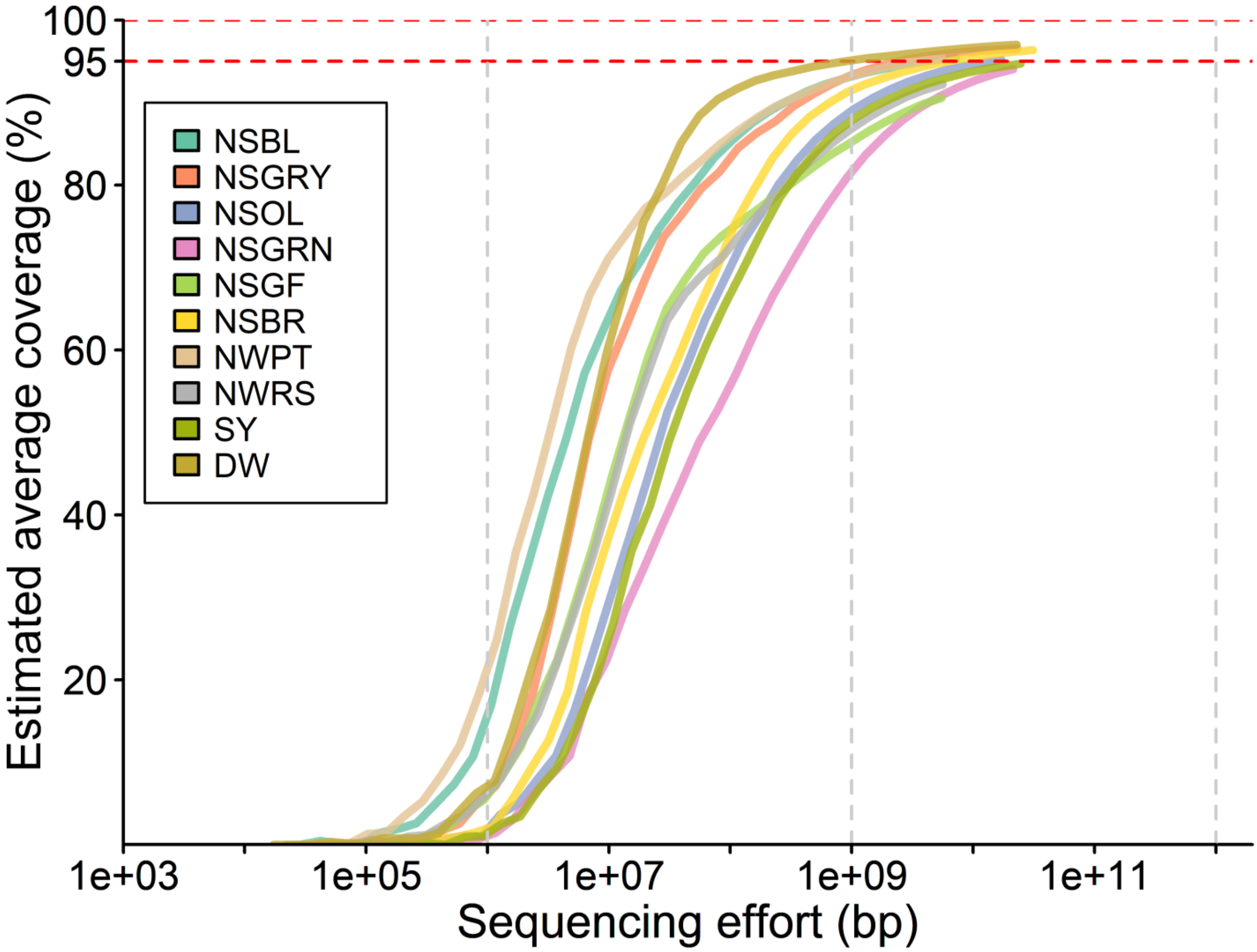


**Figure S4. Estimated metagenomic coverage of HiFi reads.**

The red horizontal dashed lines indicate 95% and 100% coverage.

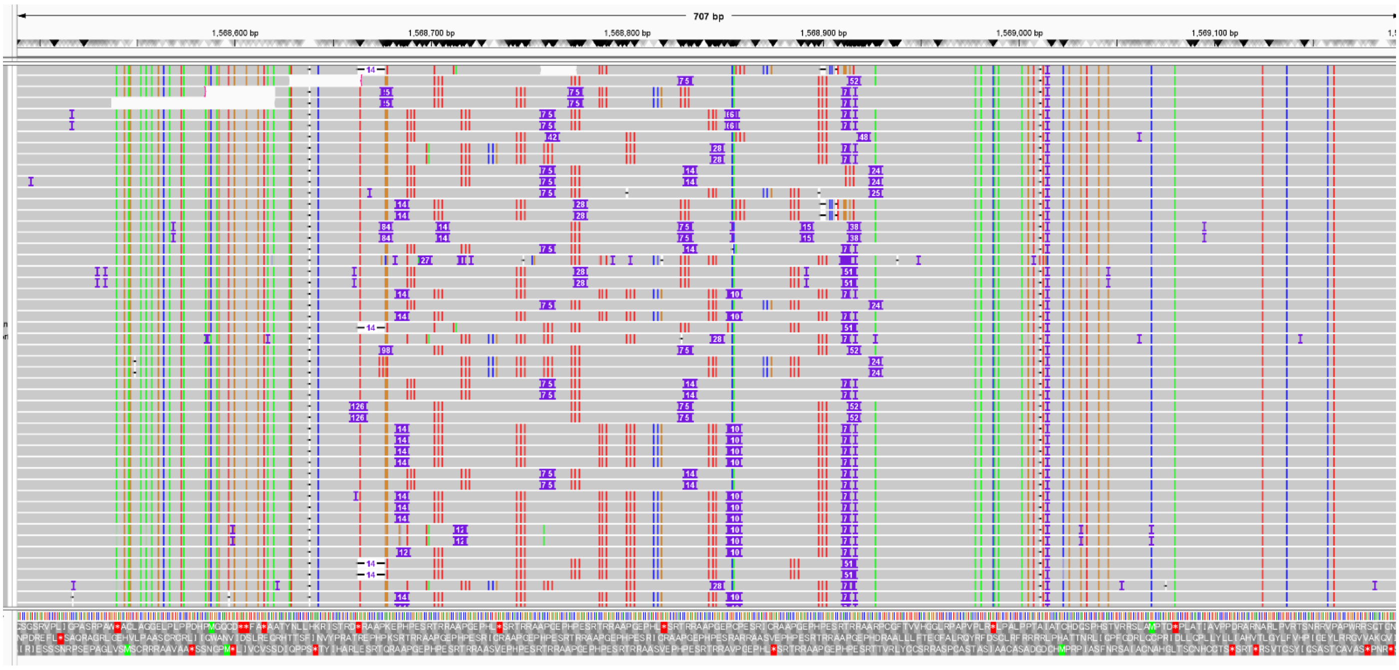


**Figure S5. HiFi reads mapped on the *Chloroflexus aggregans* DSM 9485 genome.**

As an example, a hypervariable short sequence repeat (SSR) region (1568664–1568926) is shown at base pair resolution. Each HiFi read mapped to the region shows profiles of insertions (purple box), deletions (white blank with an intermediate line), and mutations (colored stripes). The numbers in the insertion and deletion boxes represent the lengths.

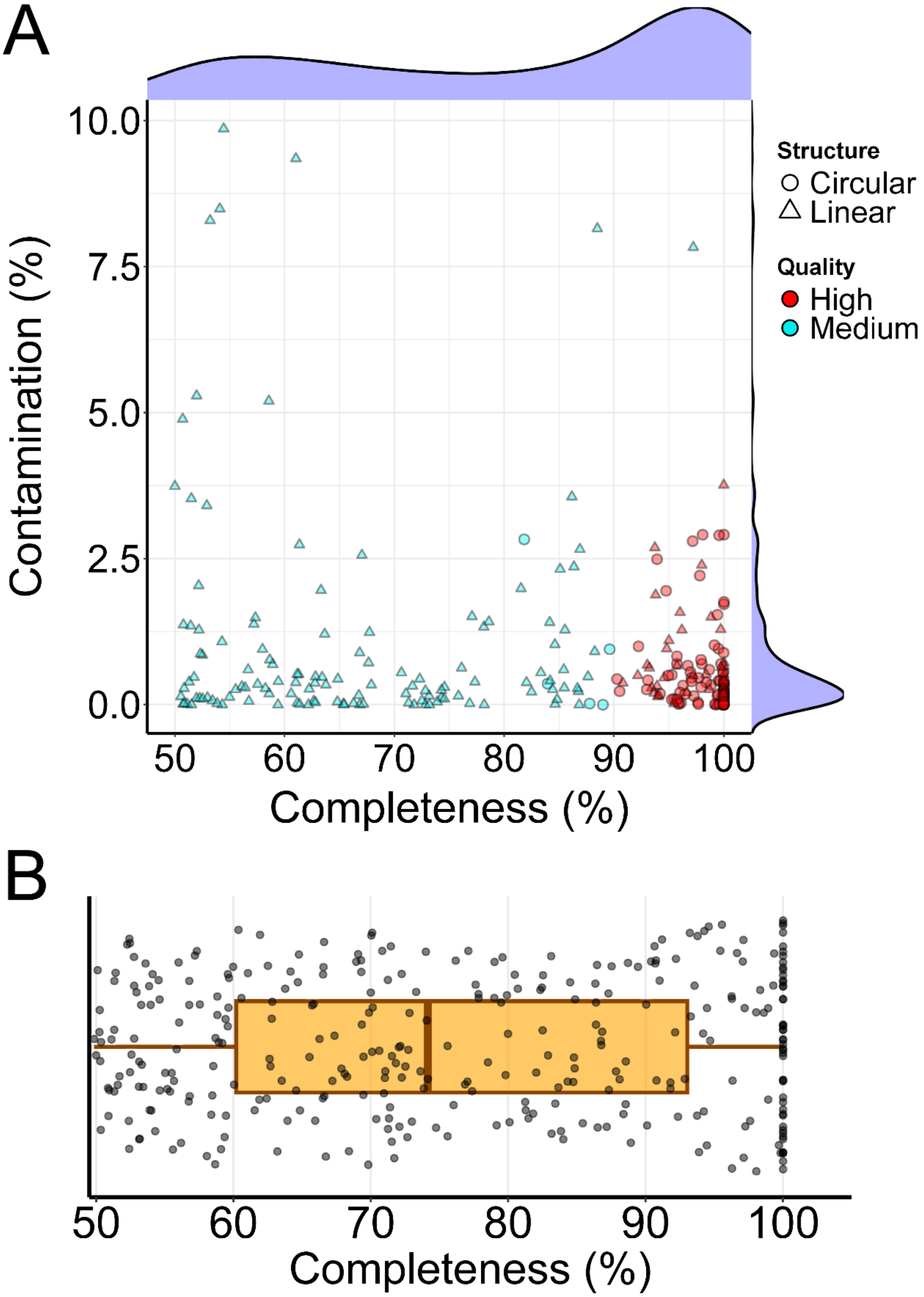


**Figure S6. Genome qualities of prokaryotes and viruses.**

(A) Completeness and contamination of pMAG/SAGs. Circular and linear genomes are indicated by a circle and a triangle, respectively. Genomes estimated to be of high or medium quality are colored red and blue, respectively. (B) Completeness of vMAGs. The solid vertical line indicates the median value, the box represents the interquartile range (25–75%), and the horizontal line indicates the 1st to the 99th percentile.

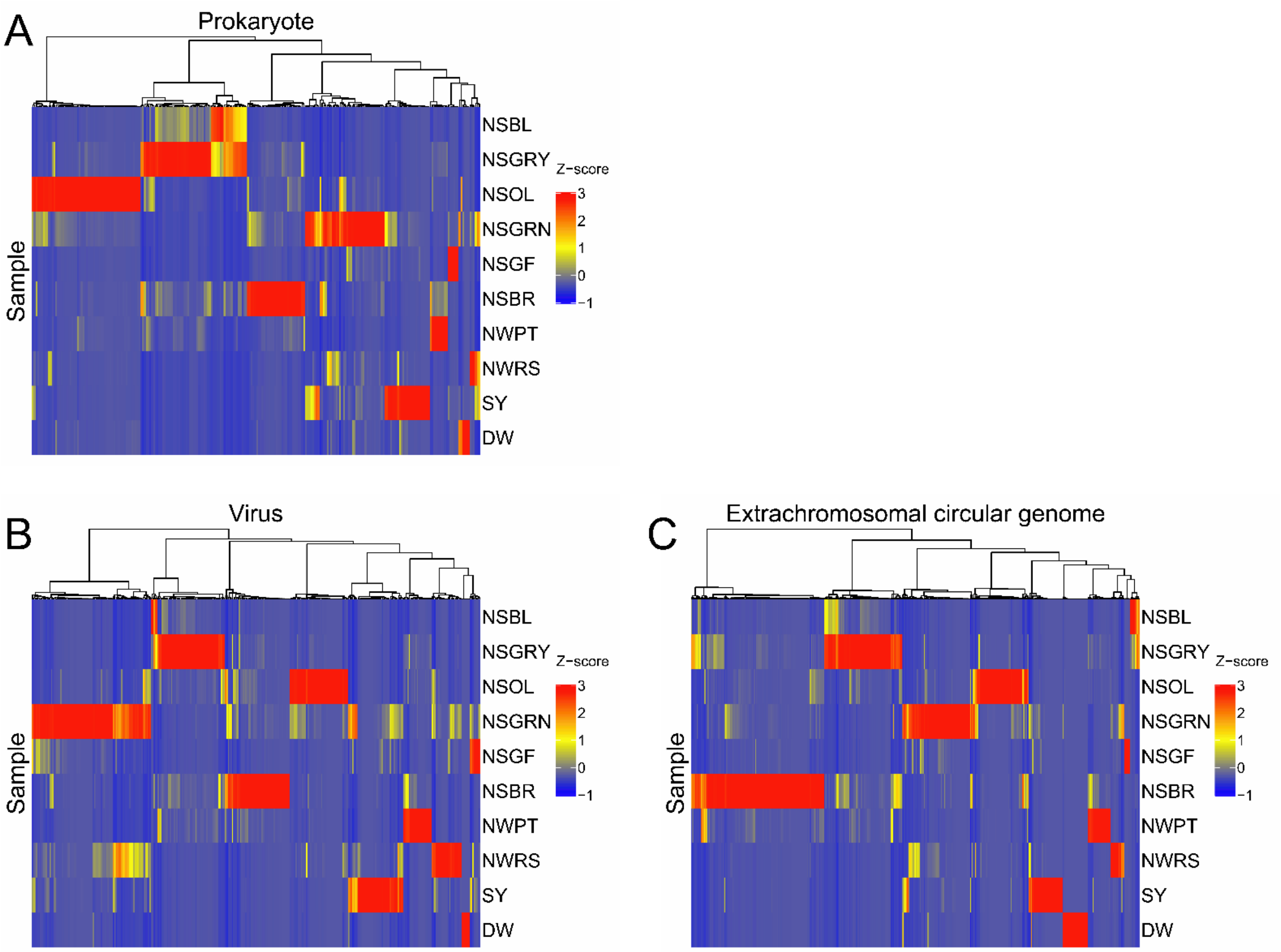


**Figure S7. Z-scored genome coverage across the samples.**

Heat map visualizing the Z-score distribution of the HiFi read coverage of (**A**) prokaryotes (pMAG/SAGs), (**B**) viruses (vMAGs), and (**C**) extrachromosomal circular genomes (eccMAGs). The dendrograms depict the hierarchical clustering of the genomes using Euclidean distance and the Ward's method.

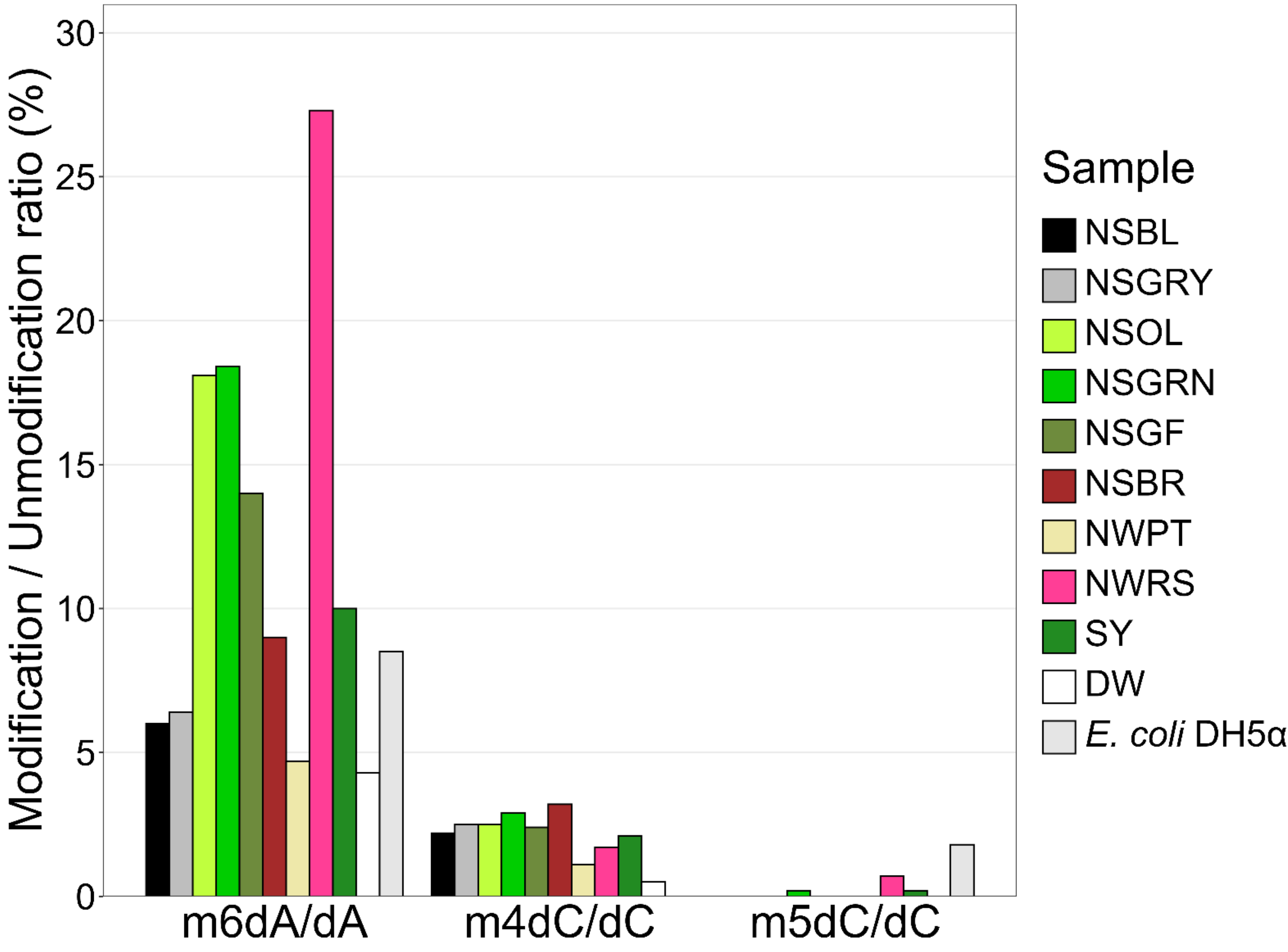


**Figure S8. Nucleosides analysis of metagenome samples**

The three methylated deoxyribonucleosides (m6dA, m4dC, and m5dC) and the corresponding unmodified deoxyribonucleosides (dA and dC) were quantified by LC-MS/MS, and the ratios were calculated. In addition to the hot spring biofilm metagenome samples, the genome of pure-cultured *E. coli* DH5α was examined.

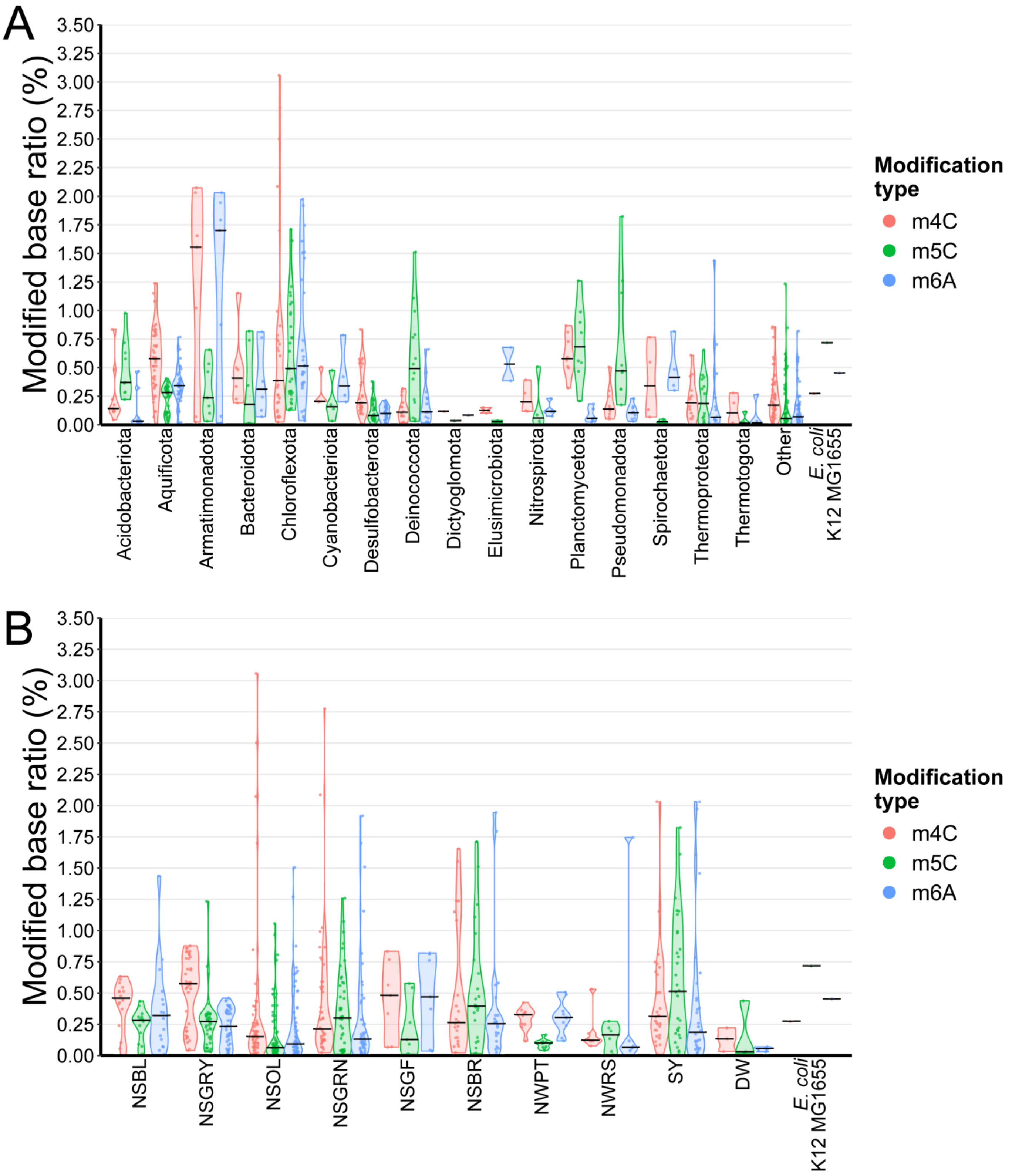


**Figure S9. Modified base ratios of the prokaryotic genomes.**

(**A**) Modification ratios by phylum. **(B)** Modification ratios by samples. In addition to the genomes produced in this study, the ratios of *E. coli* K-12 MG1655 are indicated. The solid horizontal line indicates the median value.

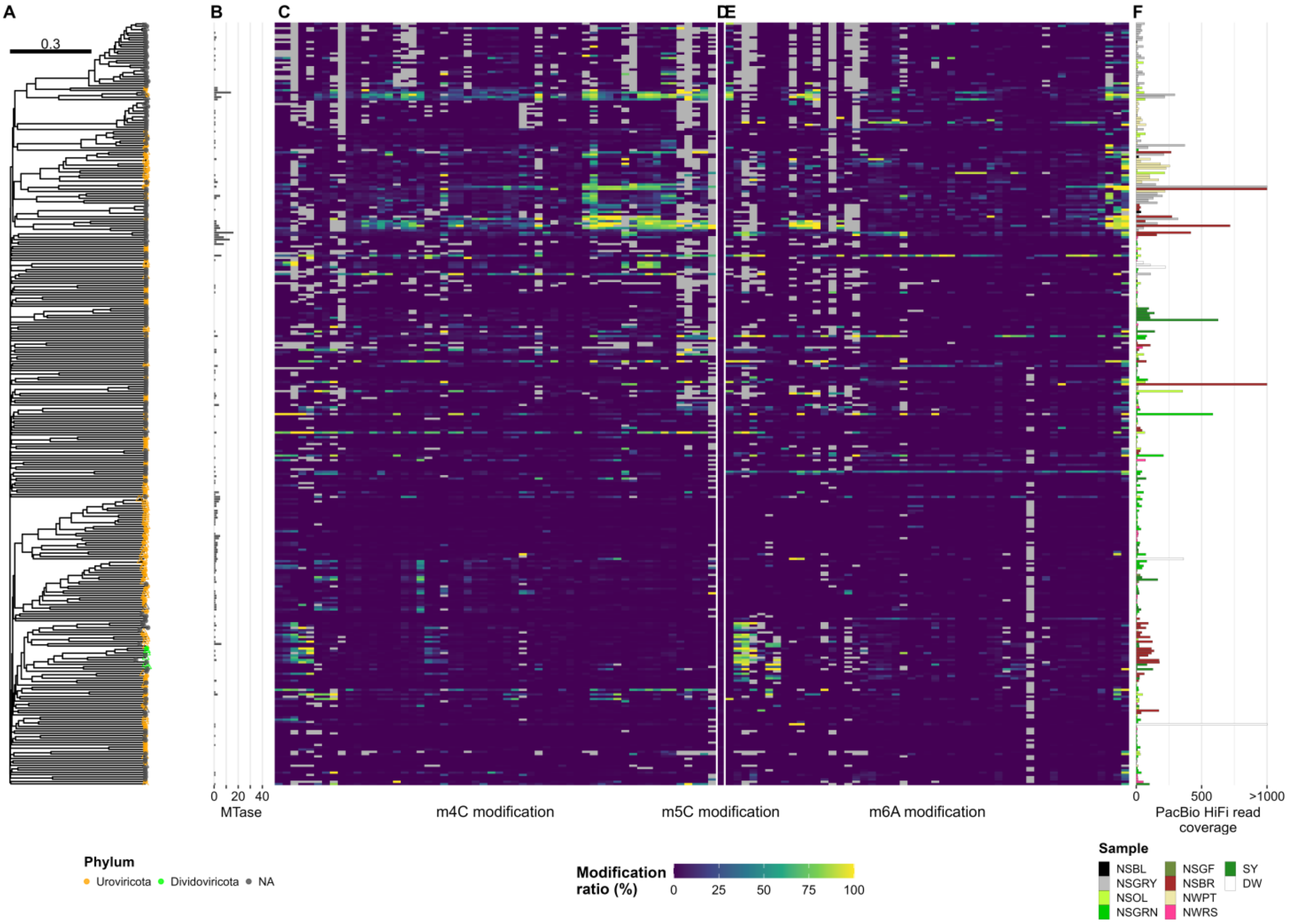


**Figure S10. Methylomes of viral genomes.**

(**A**) A proteomic tree was generated based on the vMAGs. Proviruses are indicated by a circle cross. Node color indicates taxonomy at the family level. (**B**) Number of MTase genes identified in each genome. (**C**–**E**) Modification ratios of (**C**) m4C, (**D**) m5C, and (**E**) m6A motifs. (**F**) Coverage of HiFi reads on each genome (see Figure 3).

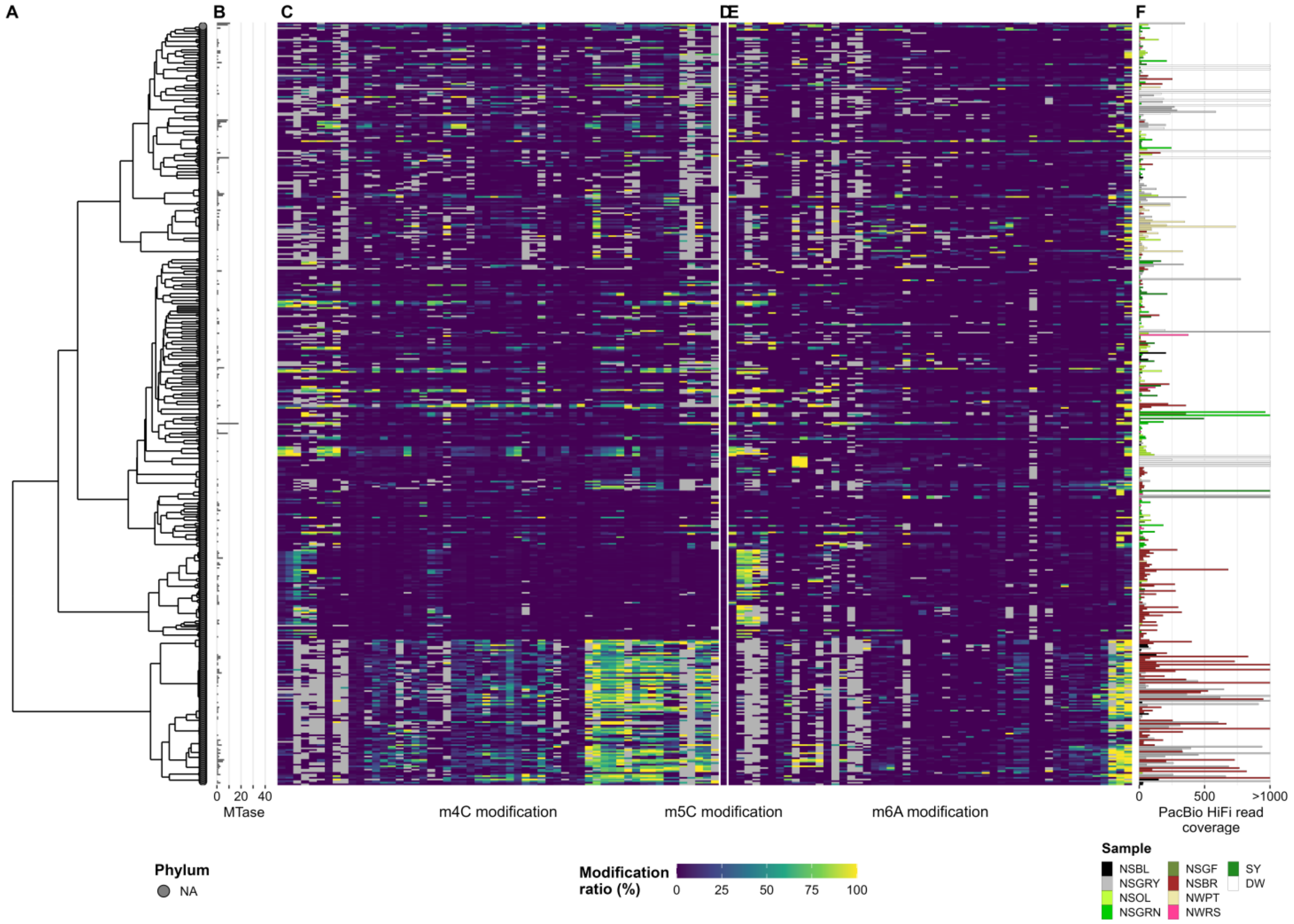


**Figure S11. Methylomes of extrachromosomal circular genomes.**

(**A**) A genome similarity dendrogram was generated based on the ANI across the eccMAGs. (**B**) Number of MTase genes identified in each genome. (**C**–**E**) Modification ratios of: (**C**) m4C, (**D**) m5C, and (**E**) m6A motifs. (**F**) Coverage of HiFi reads on each genome (see Figure 3).

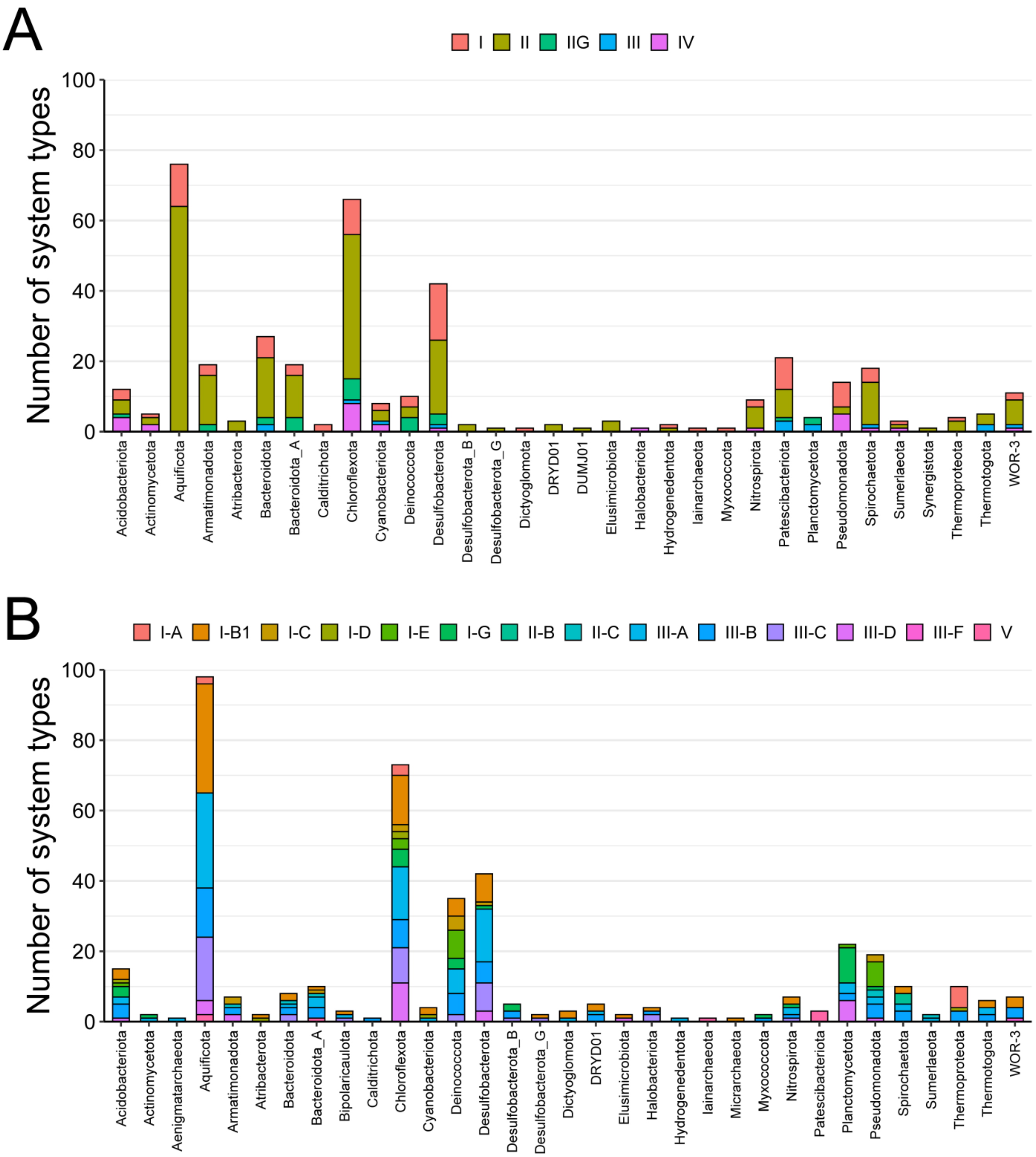


**Figure S12. RM and CRISPR-Cas systems in prokaryotic genomes.**

(**A**) RM systems for each phylum. (**B**) CRISPR system for each phylum. The bar items are colored according to the system type.

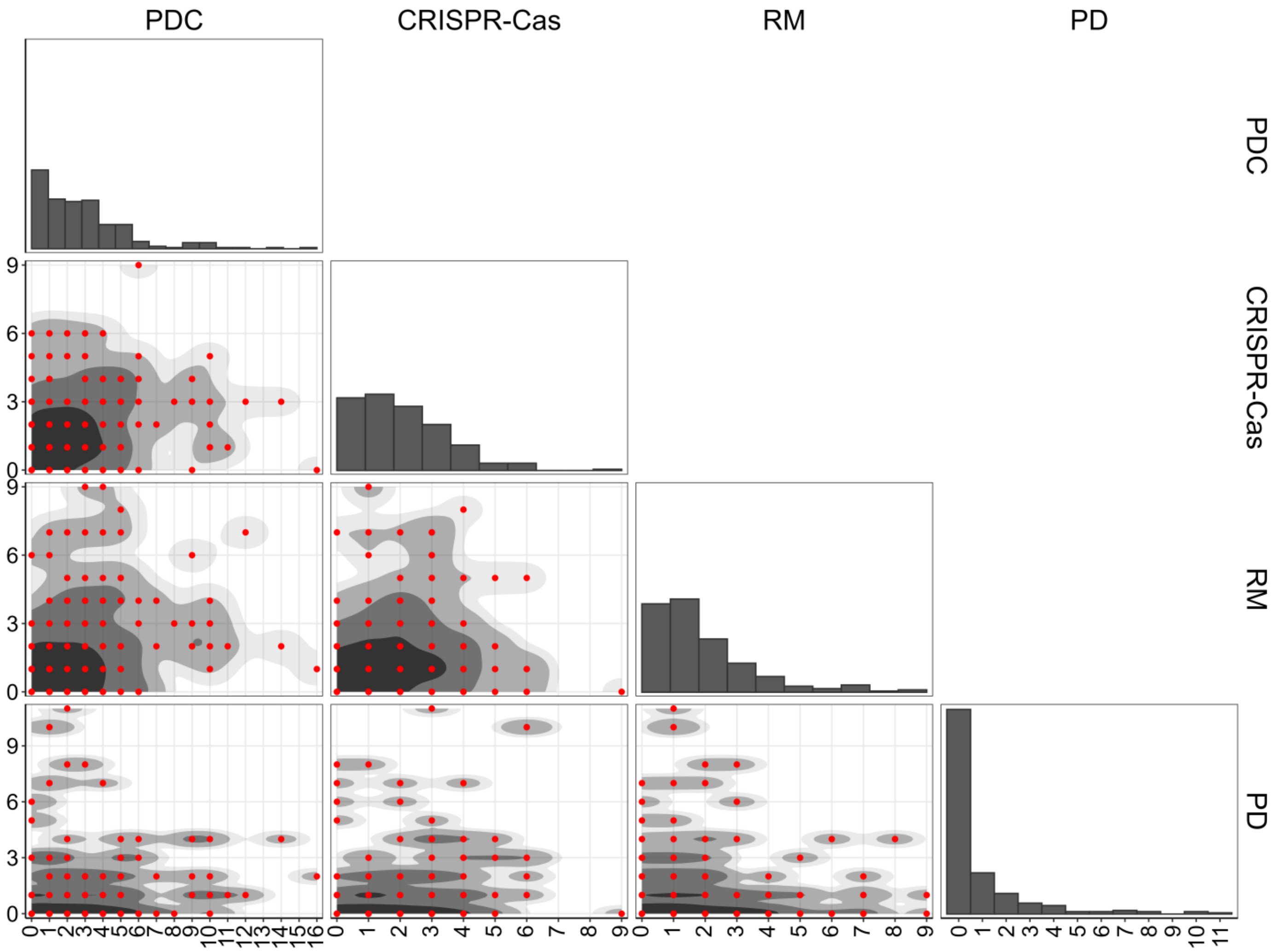


**Figure S13. Pairwise relations of PDC, CRISPR, RM, and PD systems in prokaryotic genomes.**

The lower-triangle panels show the density scatter plots for each pair of defense systems. The diagonal plots indicate the cumulative histograms of the system.

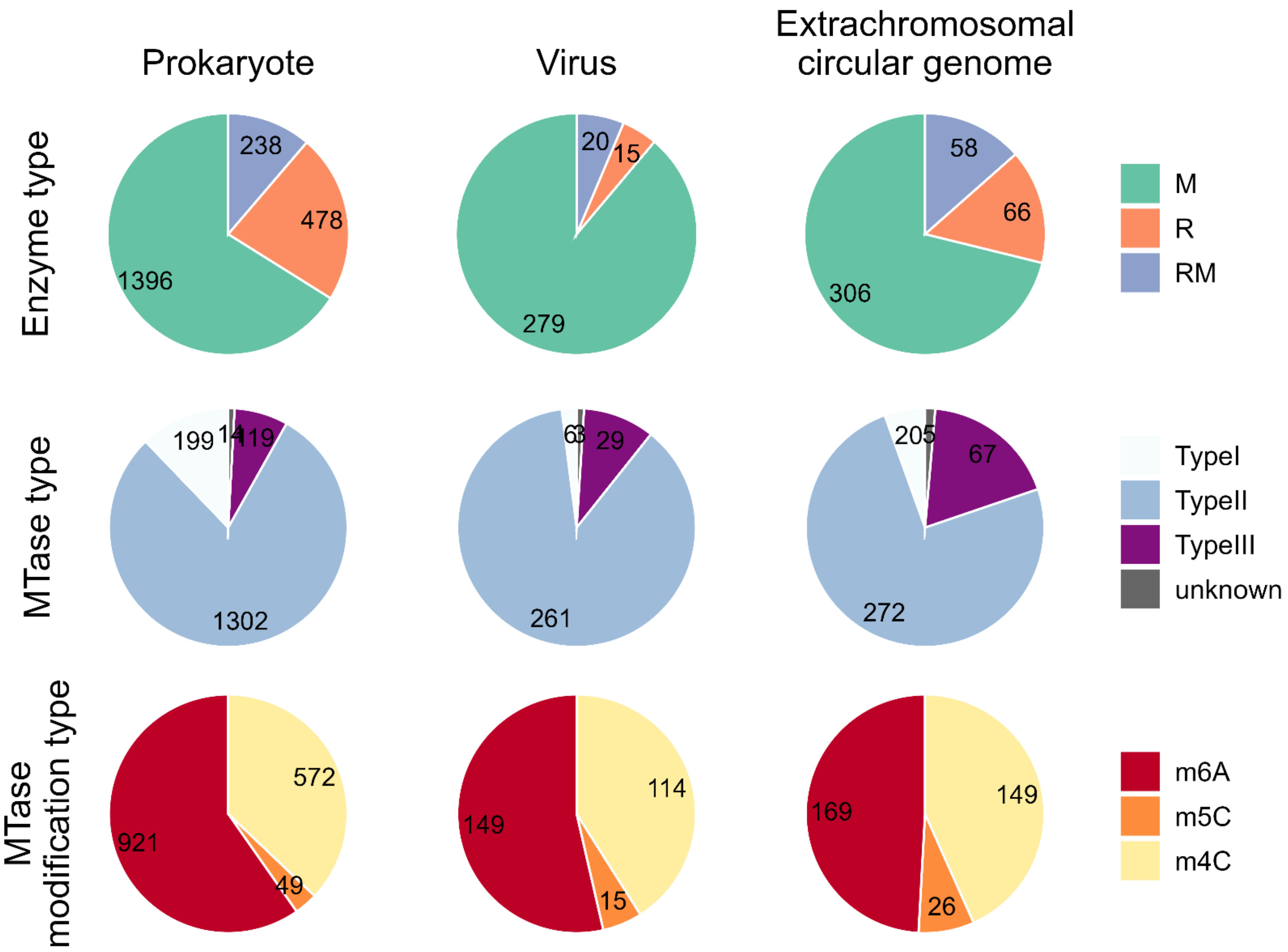


**Figure S14. Number of DNA restriction and modification system genes on the reconstructed genomes.**

Enzyme types were classified as follows: DNA methyltransferase (MTase; M), restriction endonuclease (REase; R), protein fused with M and R domains (RM), and DNA sequence-recognition protein (S).

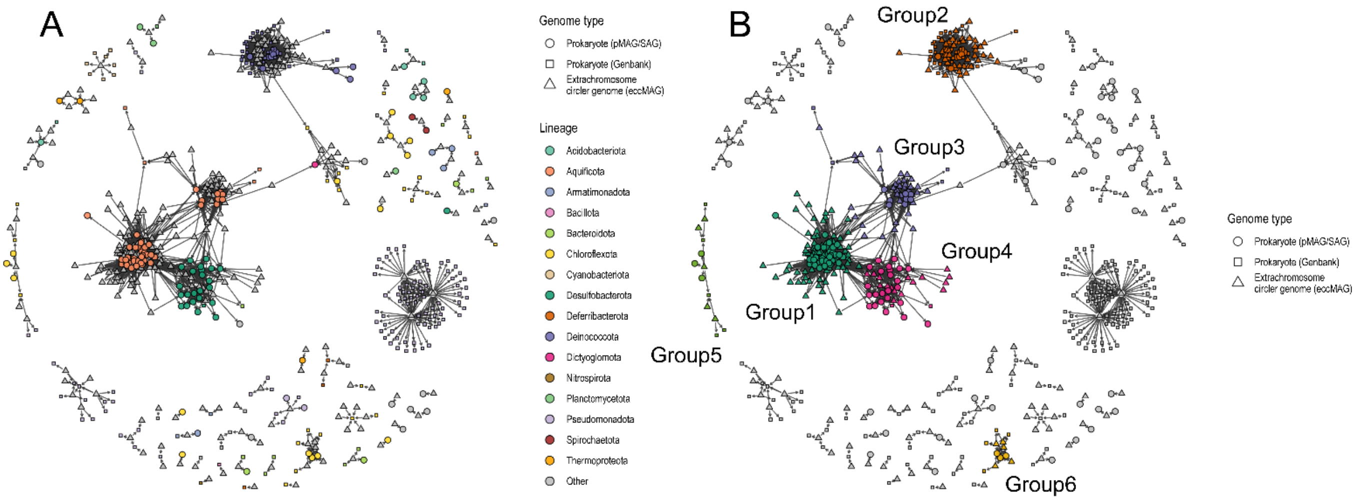


**Figure S15. Network of extrachromosomal circular genome–host interactions**

(**A**) Extrachromosomal circular genomes (eccMAGs), their estimated prokaryotic hosts (pMAG/SAGs), and public GenBank database genomes are shown as nodes in triangular, circular, and rectangular shapes, respectively. Arrows represent the interaction between host prokaryotes and eccMAGs. Major prokaryotes are colored by taxonomic assignment at the phylum level. (**B**) Network colored by subnetworks. The subnetworks were computationally determined. Nodes in the large subnetworks containing > 3 pMAG/SAGs are colored by group.

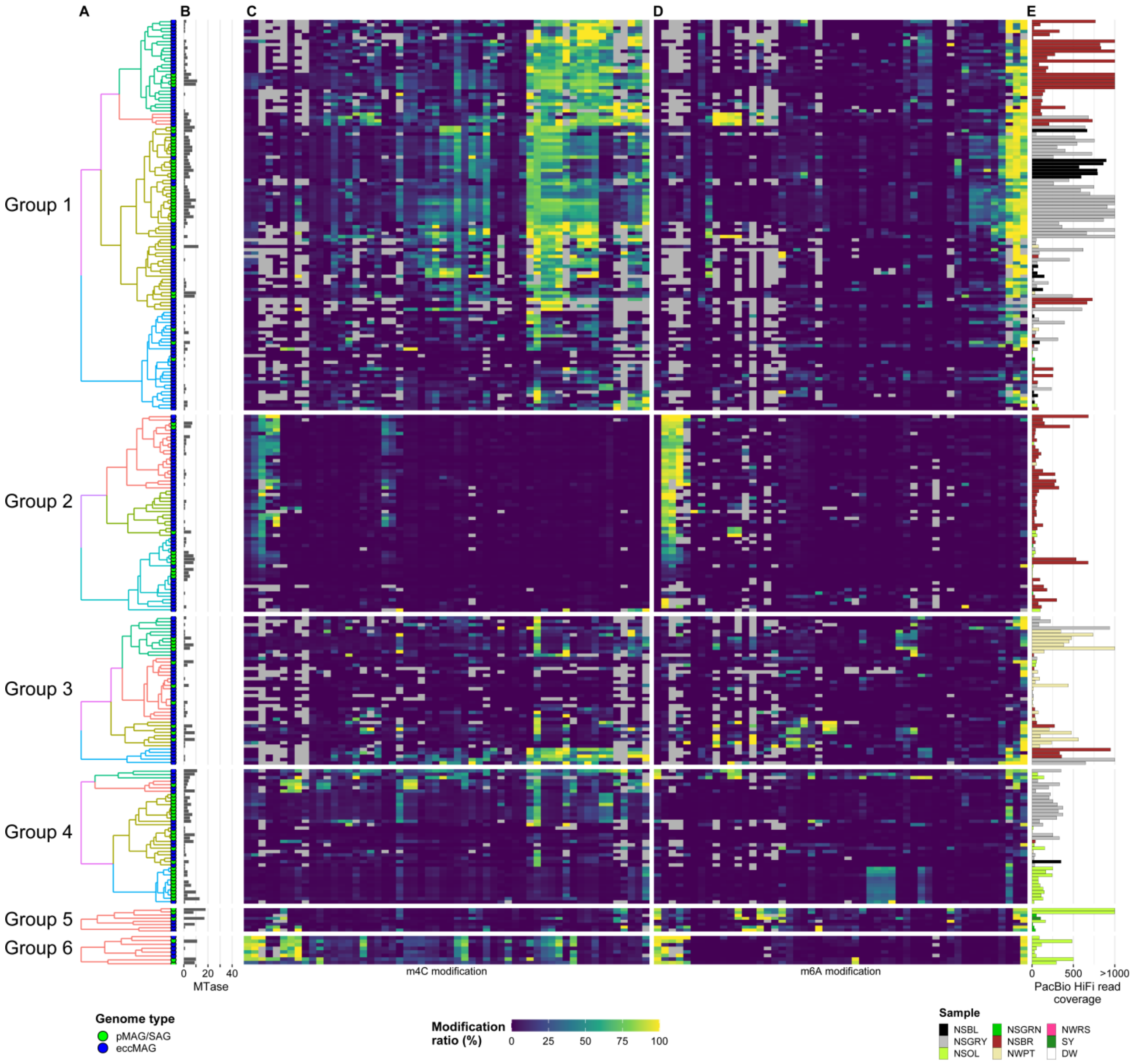


**Figure S16. Methylomes of prokaryotic and extrachromosomal circular genomes in each infection subnetwork.**

Hierarchical clustering of the modification ratio profiles within each subnetwork was performed independently using the Bray–Curtis dissimilarity index. The clusters in each group are shown in the dendrogram. Modification ratios of: (**C**) m4C and (**D**) m6A (see Figure 6).

### Supplementary Tables

#### Table S1. Description of sampling sites

| Sample | Site | Location | Sampling date | Color | Biofilm structure | Temperature (℃) | pH | Estimated dominant metabolism | Known majour members | References |
|---|---|---|---|---|---|---|---|---|---|---|
| NSBL | Nakabusa (Stream site) | 36° 23.55'N 137° 44.88'E | 2021-06-29 | Black | Streamer | 86 | 8.5–8.9 | Chemoautotroph | Thermocrinis (Aquificota)<br>Caldimicrobium (Thermodesulfobacteriota) | Kawai et al. J. Gen. Appl. Microbiol. 2025 |
| NSGRY | Nakabusa (Stream site) | 36° 23.55'N 137° 44.88'E | 2021-06-29 | Gray | Streamer | 67–77 | 8.5–8.9 | Chemoautotroph | Sulfurihydrogenibium (Aquificota)<br>Thermocrinis (Aquificota)<br>Caldimicrobium (Thermodesulfobacteriota)<br>Fervidobacterium (Thermotogaeota)<br>Thermus (Deinococcota) | Nakagawa and Fukui. Appl. Environ. Microbiol. 2003<br>Kubo et al. Syst. Appl. Microbiol. 2011<br>Nishihara et al. Microbes. Environ. 2018<br>Kawai et al. J. Gen. Appl. Microbiol. 2025 |
| NSOL | Nakabusa (Stream site) | 36° 23.55'N 137° 44.88'E | 2021-06-29 | Olive | Mat | 71 | 8.5–8.9 | Chemoautotroph (surface) – Anoxygenic phototroph (deep) | Roseiflexus (Chloroflexota)<br>Chloroflexus (Chloroflexota)<br>Caldimicrobium (Thermodesulfobacteriota)<br>Fervidobacterium (Thermotogaeota)<br>Dictyoglomus (Dictyoglomerota) | Nakagawa and Fukui. J. Gen. Appl. Microbiol 2002<br>Kubo et al. Syst. Appl. Microbiol. 2011<br>Nishida et al. Plas One. 2018<br>Hanada. Microbes. Environ. 2003<br>Everroad et. al. Microbes. Environ. 2012<br>Kawai et al. Front. Microbiol. 2018<br>Kawai et al. J. Gen. Appl. Microbiol. 2025 |
| NSGRN | Nakabusa (Stream site) | 36° 23.55'N 137° 44.88'E | 2021-06-29 | Green | Mat | 58 | 8.5–8.9 | Oxygenic phototroph (surface) – Anoxygenic phototroph (deep) | Thermosynechococcus (Cyanobacteriota)<br>Armatimonadetes bacterium (Armatimonadota)<br>Chloroflexus (Chloroflexota)<br>Chloroflexi bacterium (Chloroflexota)<br>Meiothermus (Deinococcota) | Nakagawa and Fukui. J. Gen. Appl. Microbiol 2002<br>Everroad et. al. Microbes. Environ. 2012<br>Kawai et al. J. Gen. Appl. Microbiol. 2025<br>Martinez et. al. Microbes. Environ. 2019 |
| NSGF | Nakabusa (Stream site) | 36° 23.55'N 137° 44.88'E | 2021-06-29 | Green | Mat | 51 | 8.5–8.9 | Oxygenic phototroph (surface) – Anoxygenic phototroph (deep) | Thermosynechococcus (Cyanobacteriota)<br>Chloroflexus (Chloroflexota)<br>Chloroflexi bacterium (Chloroflexota)<br>Meiothermus (Deinococcota) | Nakagawa and Fukui. J. Gen. Appl. Microbiol 2002<br>Everroad et. al. Microbes. Environ. 2012<br>Kawai et al. J. Gen. Appl. Microbiol. 2025 |
| NSBR | Nakabusa (Stream site) | 36° 23.33'N 137° 44.88'E | 2021-06-29 | Brown | Streamer | 70 | 8.5–8.9 | Chemoautotroph<br>Heterotroph | NA | NA |
| NWPT | Nakabusa (Wall site) | 36° 23.55'N 137° 44.88'E | 2021-06-29 | Pale-tan | Streamer | 75 | 8.5–8.9 | Chemoautotroph | Sulfurihydrogenibium (Aquificota)<br>Thermocrinis (Aquificota)<br>Fervidobacterium (Thermotogaeota)<br>Actinomadura (Thermotogaeota)<br>Thermus (Deinococcota) | Everroad et. al. Microbes. Environ. 2012<br>Nishihara et al. Microbes. Environ. 2018a<br>Nishihara et al. Microbes. Environ. 2018b<br>Nakagawa and Fukui. J. Gen. Appl. Microbiol 2002 |
| NWRS | Nakabusa (Wall site) | 36° 23.33'N 137° 44.88'E | 2021-06-29 | Rose | Mat | 66 | 8.5–8.9 | Anoxygenic phototroph | Roseiflexus (Chloroflexota) | Hanada. Microbes. Environ. 2003 |
| SY | Shiriyaki | 36° 38.89'N 138° 38.46'E | 2021-11-17 | Green (surface layer), Orange (deep layer) | Mat | 51 | 5.5 | Oxygenic phototroph(surface) – Anoxygenic phototroph (deep) | NA | NA |
| DW | Dewanoyu | 40° 25.71'N 140° 37.23'E | 2022-10-14 | White | Streamer | 58 | 7 | Chemoautotroph | NA | NA |

**Table S2. Water chemistry**

| Hot spring | Water sampling spot | Sampling date | pH | $Cl^-$ (mg/L) | $NO_3$ (mg/L) | $NO_2$ (mg/L) | $SO_4^{2-}$ (mg/L) | $H_2S$ (mg/L) | $PO_4$ (mg/L) | $NH_4$ (mg/L) | $Na^+$ (mg/L) | $K^+$ (mg/L) | $Mg^{2+}$ (mg/L) | Fe (mg/L) | Mn (mg/L) |
|---|---|---|---|---|---|---|---|---|---|---|---|---|---|---|---|
| Nakabusa (stream site) | N-1 | 2024-11-14 | 7.0 | 36 | NA | NA | 36 | 6.03 | NA | NA | 62 | 6.9 | 0.19 | <0.5 | <0.5 |
| Nakabusa (stream site) | N-2 | 2024-11-14 | 7.2 | 37 | NA | NA | 37 | <0.02 | NA | NA | 63 | 7.4 | 0.15 | <0.5 | <0.5 |
| Nakabusa (stream site) | N-3 | 2024-11-14 | NA | 44 | NA | NA | 39 | 5.90 | NA | NA | 73 | 9.0 | 0.06 | <0.5 | <0.5 |
| Nakabusa (stream site) | N-4 | 2024-11-14 | NA | 44 | NA | NA | 38 | 5.62 | NA | NA | 73 | 8.3 | 0.06 | <0.5 | <0.5 |
| Nakabusa (stream site) | N-5 | 2024-11-14 | NA | 44 | NA | NA | 43 | <0.02 | NA | NA | 74 | 9.2 | 0.06 | <0.5 | <0.5 |
| Nakabusa (wall site) | N-6 | 2024-11-14 | 8.7 | 51 | NA | NA | 28 | 5.39 | NA | NA | 81 | 7.4 | <0.05 | <0.5 | <0.5 |
| Shiriyaki | S | 2021-11-17 | 5.5 | 220 | <30 | <30 | 680 | 25.6 | <300 | <30 | 170 | <30 | <30 | <0.5 | <0.5 |
| Dewanoyu | D | 2022-10-14 | 7.0 | 7500 | <30 | <30 | <30 | <0.1 | <300 | <30 | 4300 | 180 | 180 | <0.5 | 4.2 |

**Table S3. Data on metagenomic shotgun sequencing read analyses and metagenomic assembly.**

| Sample | HiFi reads | HiFi read length (bp) | Max length (bp) | HiFi reads total base (bp) | Accession ID | CDSs | MTase | REase | First-step assembled contigs | Total length (bp) | N50 (bp) | Longest length (bp) | Average length (bp) | pMAG | pSAG | pMAG/SAGs (dereplicated) | vMAG | eccMAG | Mapped HiFi reads on the genome set (%) |
|---|---|---|---|---|---|---|---|---|---|---|---|---|---|---|---|---|---|---|---|
| DW | 2,974,256 | 7,667 ± 2,889 | 34,096 | 22,802,713,809 | DRR959823 | 30,328,704 | 84,630 | 28,707 | 24,668 | 427,376,828 | 18,623 | 2,044,028 | 17,325 | 3 | NA | 3 | 6 | 26 | 48.4% |
| NSBL | 599,382 | 12,017 ± 3,171 | 41,389 | 7,202,927,463 | DRR959824 | 10,728,310 | 55,246 | 22,751 | 7,066 | 294,082,748 | 45,366 | 2,971,164 | 41,619 | 25 | NA | 16 | 5 | 17 | 97.2% |
| NSBR | 2,406,900 | 12,934 ± 3,034 | 45,183 | 31,130,524,262 | DRR959825 | 44,769,923 | 266,874 | 122,383 | 32,470 | 1,150,010,208 | 37,734 | 3,978,837 | 35,418 | 38 | NA | 22 | 42 | 129 | 90.8% |
| NSGF | 482,594 | 11,394 ± 2,367 | 34,265 | 5,498,889,692 | DRR959826 | 7,199,968 | 25,955 | 12,736 | 5,829 | 228,841,328 | 47,285 | 4,961,631 | 39,259 | 7 | NA | 7 | 9 | 9 | 66.6% |
| NSGRN | 1,816,975 | 11,790 ± 2,944 | 45,861 | 21,421,664,296 | DRR959827 | 26,546,606 | 121,474 | 55,077 | 32,677 | 1,289,908,643 | 48,473 | 5,310,370 | 39,475 | 49 | NA | 44 | 80 | 69 | 89.6% |
| NSGRY | 1,887,826 | 12,294 ± 2,969 | 47,488 | 23,209,001,521 | DRR959828 | 33,026,532 | 191,618 | 80,257 | 24,478 | 879,831,992 | 38,404 | 3,090,523 | 35,944 | 58 | NA | 42 | 49 | 83 | 92.4% |
| NSOL | 1,524,084 | 11,235 ± 2,613 | 42,742 | 17,123,686,230 | DRR959829 | 21,856,311 | 82,203 | 39,875 | 21,321 | 754,463,083 | 39,934 | 5,154,045 | 35,386 | 51 | 224 | 68 | 47 | 55 | 95.2% |
| NWPT | 437,219 | 12,779 ± 2,926 | 40,270 | 5,587,035,199 | DRR959830 | 8,154,349 | 37,932 | 23,196 | 4,899 | 209,204,383 | 48,157 | 1,860,130 | 42,704 | 87 | NA | 10 | 25 | 25 | 99.2% |
| NWRS | 507,960 | 11,033 ± 2,348 | 37,510 | 5,604,137,819 | DRR959831 | 6,898,663 | 28,378 | 12,152 | 7,317 | 312,053,918 | 51,381 | 4,664,569 | 42,648 | 15 | NA | 6 | 32 | 16 | 88.7% |
| SY | 2,737,558 | 9,015 ± 4,059 | 45,831 | 24,679,702,742 | DRR959832 | 29,374,793 | 66,829 | 26,407 | 12,971 | 556,209,331 | 65,404 | 5,601,991 | 42,881 | 35 | NA | 30 | 37 | 36 | 93.8% |

### Table S4. Short sequence repeat on the *Chloroflexus aggregans* DSM 9485 genome

| NO. | Start | End | Period Size | Copy Number | Consensus Size | Percent Matches (%) | Percent Indels (%) | Alignment score | A (%) | C (%) | G (%) | T (%) | Entropy (0-2) | Period sequence |
|---|---|---|---|---|---|---|---|---|---|---|---|---|---|---|
| 1 | 95118 | 95633 | 18 | 28.8 | 18 | 86 | 0 | 568 | 12 | 44 | 34 | 9 | 1.74 | CGGGGGCCTGCCCCAACC |
| 2 | 173824 | 175083 | 37 | 33.8 | 37 | 90 | 1 | 1746 | 9 | 32 | 48 | 10 | 1.68 | AGCGCCGGGGTGCGCCGGCGGGTGACGGCCCTGCACG |
| 3 | 403711 | 405475 | 38 | 46.8 | 38 | 96 | 1 | 3252 | 9 | 41 | 33 | 15 | 1.79 | CCCCCGCCCCCGTCGGCAATCCGGTGGGTTGGCATGCG |
| 4 | 443676 | 445503 | 126 | 14.5 | 126 | 92 | 1 | 2728 | 12 | 32 | 38 | 17 | 1.86 | CAGCGCCCGCTCGTACAATGGGCGGGCGGCCGCGTAGTCGCCTTGGGATGCCAGCAGCCCGGCCAGATTGTTCAGGCTGGTGGCCGTATCGGGGTGGTCGGGCCCCAGCGCCCGTTCGCGGATTGC |
| 5 | 569984 | 573319 | 119 | 27.7 | 118 | 89 | 3 | 4533 | 21 | 30 | 30 | 17 | 1.96 | TTTGCGTCGGGGCACGGCGACGCCGTGTCGGGGCGACGCGCCGCGTCGCCCCTACGTATCAATGATGCCACGCACGTGATGGGGTATAACCCGAAATACAACCGTACATACGGACGGA |
| 6 | 662905 | 663057 | 15 | 10 | 15 | 82 | 12 | 168 | 0 | 40 | 41 | 16 | 1.54 | GGTGCCGCTGCTGCC |
| 7 | 822885 | 827638 | 306 | 15.2 | 306 | 75 | 11 | 2876 | 16 | 28 | 40 | 14 | 1.87 | ACGGCCATTGCAGCGGGTGACGATCACACCTGTGCGCTGACGAGCAGCGGCGGGGTGTGGTGCTGGGGATACAACCTCTACGGGCAACTGGGCGACGGGACGACGACCAACCGCAGCACGCCGGGGGCGGTCAGCGGGCTACCGAGCGGGGTGACGGCCATTGCAGCGGGTGATGATCACACCTGTGCGCTGACCGGCAGCGGCGGGGTGTGGTGCTGGGGCGACAACGACTTCGGGCAACTGGGCGACGGCTCGTACACCGACCGCAGCACGCCGGGGGCGGTCAGCGGGCTACCGAGCGGGGTG |
| 8 | 838467 | 840715 | 107 | 20.3 | 107 | 83 | 9 | 2337 | 21 | 31 | 28 | 19 | 1.97 | TGCGAGCGCAGCCTGTCCTTGCGAGCGTAGCCTGTCCTTGCGAGCGCAGCGAGAAATCTTGTCAAGATTGCTCAGTCGCTGACGCGCCCTCGCAATGACAAAGGAAC |
| 9 | 897528 | 899821 | 99 | 21.9 | 99 | 84 | 10 | 1903 | 16 | 45 | 28 | 10 | 1.8 | GCCCGCCACGCCGCCGACGCGATTGCTGACAACGATGACCGCGCCAAGGCGCTGACCGAACTTGCCACCGCCCTCGCCCACGCCGGGCAGTTCACCGCC |
| 10 | 1005974 | 1006095 | 12 | 10.2 | 12 | 97 | 0 | 226 | 41 | 56 | 0 | 1 | 1.09 | ACCAACACACCC |
| 11 | 1007648 | 1009050 | 72 | 19.4 | 72 | 76 | 15 | 1138 | 29 | 48 | 20 | 2 | 1.62 | ACGCCCACCAACACGCCGACGAACACGCCAACGAACACGCCCACTTACACGCCGACGCCGACCGAGACGCCG |
| 12 | 1065795 | 1067758 | 190 | 10.1 | 189 | 92 | 3 | 2901 | 15 | 28 | 42 | 14 | 1.85 | ATGTGGGGGACGGCATGTCGGGGCGACCCGCCGGGTCGCCCCTACAATGGGCGACGCACCGCGTCGCCCCGACGCGAACCGGCGCGGCATGTGGGGCCACGGTACGGCATGTATGTAGGGGCACGGCATGCCGTGCCCAACGGGGGTTACGTTGTTGCGGGGTGGGGGGACGGCATATCGGGGCACGGC |
| 13 | 1097936 | 1099350 | 100 | 14 | 101 | 80 | 10 | 1420 | 16 | 48 | 24 | 10 | 1.78 | CCCACCCGCACGGCATTGCCGTGCCCCGACGCCACCCACACGCACGGCATTGCCGTGCCCCGACGCCACCCACACGCACGGCATTGCCGTGCCCCGACGCA |
| 14 | 1152615 | 1152871 | 10 | 25.2 | 10 | 96 | 3 | 460 | 0 | 51 | 38 | 9 | 1.34 | CCGTGCGGCC |
| 15 | 1280069 | 1281457 | 54 | 25.5 | 54 | 70 | 12 | 345 | 1 | 31 | 41 | 25 | 1.66 | GTGGCCGTCGCCGTCGCCGTCGGCGTGGCCGTTGCCGTCGGCGTGTCCGTTGGC |
| 16 | 1305211 | 1306146 | 54 | 17.3 | 54 | 99 | 0 | 1863 | 14 | 31 | 35 | 18 | 1.91 | GCCGCACAGCCGTGCGGCGCTGCCCATTGATCGCGATCACCGTATGGTTGGGGA |
| 17 | 1430216 | 1430433 | 9 | 22.2 | 9 | 62 | 23 | 50 | 5 | 33 | 22 | 38 | 1.77 | TCCTTTGCG |
| 18 | 1442525 | 1443421 | 60 | 14.9 | 60 | 95 | 1 | 1575 | 23 | 32 | 27 | 16 | 1.96 | GTCATTTCGCGGGAGCGAAGCGACCGAGAAATCCTAACGCCAAGTCGCTACGCTCGCTCG |
| 19 | 1513072 | 1513741 | 40 | 16.7 | 40 | 92 | 1 | 1038 | 7 | 43 | 22 | 26 | 1.8 | GCGTCTGTGCGCACTTCGCGCCTTTGCGTTACCCCCCTCA |
| 20 | 1568664 | 1568926 | 14 | 18.8 | 14 | 89 | 0 | 409 | 22 | 39 | 34 | 3 | 1.72 | GAGAGCCGCACCCG |
| 21 | 1604158 | 1604651 | 34 | 14.5 | 34 | 97 | 0 | 934 | 11 | 40 | 38 | 8 | 1.72 | GCCGCACGGCAGCGCGGCTCATGCGGCGCATCCG |
| 22 | 1613418 | 1614814 | 21 | 66.5 | 21 | 94 | 0 | 2326 | 13 | 47 | 23 | 15 | 1.8 | CCACCGGCTCGGCCTCATCGA |
| 23 | 1643194 | 1643946 | 19 | 39.3 | 19 | 83 | 6 | 880 | 18 | 39 | 34 | 6 | 1.77 | ACGCACCACCGCAGTGGCG |
| 24 | 1752737 | 1754212 | 57 | 26 | 57 | 98 | 0 | 2638 | 23 | 36 | 35 | 5 | 1.79 | GACGCCGAACGACGACCATGACGCCGGACGACGGTCCCCGTAGAGACGCACGGCCGG |
| 25 | 1763007 | 1763666 | 48 | 13.9 | 48 | 95 | 2 | 1192 | 6 | 36 | 33 | 23 | 1.82 | TGCGGACGCTGAGCCGCCCTGCCGTGCGGCTTTACCGTCTTGCGCTCG |
| 26 | 1819562 | 1819929 | 25 | 14.7 | 25 | 97 | 0 | 646 | 13 | 36 | 42 | 7 | 1.73 | GCGCACGGCAGTGCGCCCGGTAGAC |
| 27 | 1878345 | 1878439 | 9 | 10.8 | 9 | 70 | 22 | 60 | 36 | 22 | 38 | 2 | 1.66 | CGCAAAGGG |
| 28 | 1922568 | 1923693 | 98 | 11.5 | 97 | 90 | 3 | 1579 | 21 | 34 | 28 | 15 | 1.94 | GACTATCGCCCCACCTCAGACGCACTGATGTCGGTATGTGCGCTGAACAGGCAAGCCACAAGCGGGTCGCACCGCGCTACCCGGTGCATTCGCCAGG |
| 29 | 1998728 | 1999665 | 56 | 17.1 | 56 | 67 | 14 | 328 | 19 | 34 | 28 | 17 | 1.95 | AGCGCAGCACCGGCCTCCTGTCATTTCAAGCGCAGCACAGGTCTCCTGTCATTTCG |
| 30 | 2096711 | 2097216 | 36 | 14.1 | 36 | 99 | 0 | 985 | 13 | 46 | 33 | 6 | 1.68 | CGCCGGGAACCCACCGCCGCCATTGGGCCGCACGGC |
| 31 | 2114118 | 2114607 | 42 | 10.9 | 42 | 68 | 21 | 230 | 14 | 55 | 24 | 5 | 1.61 | CCGCAGCCCACCGCCACCGGCCTACCGCAGCACCGCGCCGCC |
| 32 | 2207511 | 2207651 | 8 | 17.9 | 8 | 71 | 13 | 87 | 40 | 22 | 34 | 2 | 1.69 | CGCAAAGG |
| 33 | 2222211 | 2224661 | 126 | 19.5 | 126 | 88 | 4 | 2618 | 16 | 45 | 24 | 13 | 1.83 | TCCGGGCTCACCGCCACCGCAGTCACCACACTCGTATGCCCCTCCAGGGAGCGCAGCAGGCGCCCCGTCGCCGCCTCCCACACCTTCACCGTGCGATCCCAGGAACCGCTGACGATCCAGCCGCCG |
| 34 | 2304855 | 2305135 | 24 | 11.7 | 24 | 98 | 0 | 508 | 8 | 30 | 41 | 19 | 1.8 | ACCGGCGTGTTGGTCGGCGGCACC |
| 35 | 2439624 | 2439760 | 12 | 11.4 | 12 | 79 | 0 | 67 | 0 | 32 | 42 | 24 | 1.6 | GCCGTCGCCGTC |
| 36 | 2481711 | 2481793 | 8 | 10.2 | 8 | 73 | 17 | 73 | 36 | 22 | 38 | 2 | 1.68 | CGCAAAGG |
| 37 | 2621839 | 2622083 | 21 | 11.7 | 21 | 94 | 0 | 409 | 13 | 51 | 33 | 1 | 1.49 | GCCCCGCACCGCCGGGGAACC |
| 38 | 2770696 | 2771531 | 39 | 20.2 | 39 | 86 | 5 | 799 | 16 | 35 | 37 | 11 | 1.85 | CAACGGGCGGTCATCTGACCACCGCACGGCACCGGTCGG |
| 39 | 2819870 | 2820214 | 21 | 16.5 | 20 | 74 | 7 | 240 | 30 | 15 | 41 | 12 | 1.85 | ATGGGAAGAGCAAATCGGCG |
| 40 | 2825259 | 2825454 | 19 | 10.3 | 19 | 99 | 0 | 383 | 10 | 41 | 37 | 10 | 1.75 | CGGCAGTGCGGCGCCCATC |
| 41 | 2910166 | 2910777 | 16 | 37.7 | 16 | 79 | 2 | 614 | 2 | 31 | 51 | 15 | 1.56 | GCCGTGGGGCCGGTGC |
| 42 | 2912830 | 2913287 | 23 | 19.8 | 23 | 94 | 1 | 729 | 7 | 30 | 53 | 8 | 1.59 | GAGCCGCACTGCCGTGCGGGGGG |
| 43 | 3169100 | 3169431 | 31 | 10.7 | 31 | 97 | 0 | 619 | 18 | 27 | 41 | 11 | 1.86 | GAGCCGCACGGCAGTGGGGCTCAGATCAGTG |
| 44 | 3245766 | 3245860 | 9 | 10.8 | 9 | 67 | 17 | 56 | 21 | 15 | 51 | 11 | 1.75 | GACGGATGG |
| 45 | 3268990 | 3269549 | 36 | 16 | 36 | 85 | 7 | 787 | 22 | 25 | 32 | 20 | 1.98 | AGATGTCTCGCTTCGCTCGACATGACACGGAAGGGA |
| 46 | 3279657 | 3280398 | 30 | 24.9 | 30 | 98 | 0 | 1325 | 0 | 36 | 42 | 20 | 1.59 | GGCGGTGCGCCTTGCTGTGCCGGTGCGCCC |
| 47 | 3328601 | 3329227 | 30 | 21.3 | 30 | 72 | 17 | 433 | 0 | 15 | 53 | 30 | 1.43 | GTCGCCGTGGGCGTGTTGGTGGGCGTGTTG |
| 48 | 3373843 | 3374854 | 92 | 11 | 92 | 99 | 0 | 1979 | 20 | 23 | 29 | 26 | 1.99 | ATCAGGACACGGGTTCAGACGGATCTAACGGATTTGCACGGTGTGCGCGATCCGTGGTCATCTGGCAAACTCGTGTTTATCCGTGTGCTAAT |
| 49 | 3400169 | 3400555 | 18 | 21.5 | 18 | 95 | 0 | 666 | 14 | 32 | 41 | 11 | 1.82 | TGCCGTGCGGAGACGCAC |
| 50 | 3452214 | 3453520 | 111 | 11.8 | 111 | 97 | 0 | 2398 | 17 | 30 | 40 | 10 | 1.84 | GAGCCGGAGCTGCAACCGTTCTCGCTGGAAGAGCTGGGGCTGTCGCCGGAGGAGATAGCACAGCTTGAGGCGGCGCAGCGCGGGGAGCCGGCAGCCCCGGCAGCCCCGGCC |
| 51 | 3512350 | 3512974 | 40 | 15.9 | 40 | 80 | 8 | 678 | 6 | 23 | 52 | 17 | 1.67 | TGGGGCGGTGGTGCCGGGGCACGGCATCGCCGTGCGGGTG |
| 52 | 3538220 | 3538524 | 24 | 12.5 | 24 | 95 | 3 | 511 | 8 | 24 | 56 | 10 | 1.61 | GCGAGCCGCACCGGTGGGGGGTGG |
| 53 | 3608637 | 3608737 | 8 | 13 | 8 | 73 | 14 | 68 | 34 | 21 | 40 | 2 | 1.69 | CGCAAAGG |
| 54 | 4022796 | 4023332 | 36 | 15 | 36 | 98 | 0 | 1042 | 11 | 33 | 41 | 13 | 1.81 | GTAGCGCCGGACGGCTGTCCGGTTCGCCAAGGCGCG |
| 55 | 4079543 | 4080259 | 72 | 10 | 72 | 97 | 0 | 1353 | 13 | 42 | 29 | 13 | 1.83 | CATCACCGGCCATATCCCACCCGGTAGGGGCGACCCGGTGGGTCGCCCCATCGCCCCGATGGGTCGCCCACG |
| 56 | 4153028 | 4153579 | 14 | 40.3 | 14 | 71 | 14 | 267 | 17 | 38 | 39 | 5 | 1.72 | GCCGGAGCCGCACA |
| 57 | 4295664 | 4296762 | 80 | 13.8 | 80 | 93 | 2 | 1856 | 22 | 35 | 34 | 7 | 1.83 | CGCCCCCCGTAGGGGCACGGCGGTACCGTGCCCACGTCCCCACGAAGGGCACAAGGGCGCAGAGGGGTTAACGCAACAAG |
| 58 | 4356884 | 4358762 | 134 | 13.9 | 134 | 91 | 3 | 2934 | 20 | 19 | 42 | 17 | 1.89 | GGATGAGTGAGGGGACACGGATGGGACGGATGGGACGGATGAGCACGGATCGGAGGGTGTGATCCGTGTTAATCCGTTCAATCCGGCTCATCCGTGTGCGCTGGCCGTGAGGGGAGACGGATGGGAGGGGACAC |
| 59 | 4393228 | 4394551 | 125 | 10.6 | 125 | 97 | 0 | 2468 | 15 | 20 | 44 | 20 | 1.87 | GGTACCGTGGGTAGTAGGGGCACAGCGGTGTTGTGCCCCTACAGGTGATGGGGTAGGGTGTGGGTGCGGGCGGGCACGGCTGCTGTGGTGGTATTAATGATACGATGCACGCCATCACCGGGTGG |
| 60 | 4408486 | 4409063 | 17 | 34.3 | 17 | 86 | 3 | 749 | 13 | 51 | 28 | 6 | 1.66 | CGGCGTGCAGCCCCACC |
| 61 | 4434636 | 4436485 | 96 | 19.1 | 96 | 85 | 5 | 2393 | 16 | 30 | 43 | 9 | 1.8 | CCGGCGTGGTTGCGGGCGGAAGAGCCGACGGCGCCGACCAGCGACGTACCGGCGTGGTTGCGGGCGGAGGAGCCGACGGCGCCGACCAGCGACGTA |
| 62 | 4489343 | 4491125 | 120 | 14.9 | 120 | 95 | 1 | 2974 | 13 | 35 | 34 | 16 | 1.88 | AGCGTGGCGCTCTTGCCTTGCACATCGCCCAGGCGTTCCTGGATGGCGAGGGACTGCTCGTAGAGCCGCAGCGCCCCGTCGAGGTCGCCGCGCGTCACCAGCACGCCGGCCATGTTGTGG |
| 63 | 4555092 | 4555747 | 13 | 50.5 | 13 | 97 | 0 | 1078 | 19 | 46 | 34 | 0 | 1.52 | CCGGGCCGCACAA |
| 64 | 4581985 | 4582226 | 24 | 10.5 | 24 | 90 | 5 | 376 | 1 | 24 | 60 | 13 | 1.4 | GGCGTGGCCGTGGGTGGCGGCGGC |
| 65 | 4648785 | 4649115 | 30 | 11 | 30 | 96 | 0 | 599 | 12 | 40 | 37 | 9 | 1.76 | CCTGCCGGAGCCGTCGGAGCCTGAGCCGCG |
| 66 | 4654267 | 4654775 | 9 | 56.6 | 9 | 85 | 5 | 654 | 0 | 53 | 31 | 14 | 1.47 | CGGCTGCCC |
| 67 | 4670456 | 4670948 | 19 | 25.8 | 19 | 97 | 2 | 927 | 5 | 63 | 25 | 5 | 1.38 | CGGCCGTGCGCCCCCACCC |

#### Table S5. Novel MTases whose specificities were experimentally assayed.

| Gene ID | Estimated motif | Gene length (aa) | Gene type | RM type | MTase assay: in vitro one-pot approach | MTase assay: in vivo heterologous expression in *E. coli* | MTase assay: in vitro purified enzyme assay | Amino acid sequence of MTase | Extra amino acids at C-terminus for expression vector | Used REases for digestion assay | note |
|---|---|---|---|---|---|---|---|---|---|---|---|
| pMAG_1st_NSGRY_Circle_017_413 | AGCT(3,m4C) | 275 | M | II | Negative | Negative | NA | MPPQDMKNLFLNKIICGDVLEVLSQIPSDSIDLGITLPPYNNKKERYGGWLVDKVIYKGAKDKMNEEEYQNWQIEVLNELYRITKDGGSFFYNHKVRYEDGKMLHPVSWLLKTKWNVWQEIIWYRKIAGNIRGWRFWQVEERIYWLVKGKPKELKPEHAKFTSVWEIRPESGHKEHPAVFPIELPARIIYSILDQKGIVIDPFCGTGTTCVAAKLLGHDYIGIDISPEYVDYATKRIENAEKEKLRVISEISQHSVELTFEERKSIGLWDKKIKK | NA | NdeI (linearize), AluI | |
| pMAG_1st_NSBR_Circle_009_1219 | CAGNNNNNNTAYC(2,m6A) GRTANNNNNNCTG(4,m6A) | 301 | M | I | Negative | NA | NA | MMQNCGAWRMPWGDSMNAAGYKHVVLALIFLKYISDAFEEHRKRLEHESYADLEDPDEYRGQNVFWMPPKARWECLQKDWKFGVPPLRNANFAWGQHIIHHLAPTGYAGFVLANGSMSSNQSGEGEIRKNIIEADLVDCMVALPDKLFYSTQIPACLWFLARDKSGRPPTGRKEPLRDRRGEILFIDARRMGKMVDRIHRELTDEEIQRIARTYHAWRGEKEAGAYQNVPGFCKSTTLDEVRKHGYVLTPGRYVGAEVQQDDEPFEEKMARLVAELREQQAEAARLDEAIWKNLKALGYDA | NA | NA | |
| pMAG_1st_NSBR_Circle_009_1222 | CAGNNNNNNTAYC(2,m6A) GRTANNNNNNCTG(4,m6A) | 442 | S | I | Negative | NA | NA | MAGEWVKRKIGSLGRVVTGKTPTTADSSNFGGPYPFITIPDLDGRVLIDRTERTLSERGAEILRSCLLPPGAVMMSCIATVGKCGITTQPSFTNQQINSVIPNDDVDSRFLYYVFTQLGHELESSGSGGSVYTNVSKSRFSNIEVLIPVDLGEQRAIAHILGTLDDKIELNRRLSETLEQMARALFKAWFVDFEPVRAKMEGRWQRGQSLPGLPAHLYDLFPDRLVESELGEIPEGWEVKAIGEMAEVVGGSTPKTNRAEFWEGGTYHWVTPKDLSHLSLPVLLDTERKITDAGLAQISSGLLPQGTVLLSSRAPIGYLAIAEVPVAVNQGFIAMRPRQGVSNLFLLHWARAAHDEIISHANGSTFLEISKASFRPIRLVAPPPAVMLAYDRLSRPLYRRVVENERESHTLAALRDALLPKLIRGEIRVKDAERFLKERGLC | NA | NA | |
| pMAG_1st_NSGRY_Circle_004_1602 | CCGG(2,m4C) | 274 | M | II | Negative | NA | NA | MPPQDMKNLFLNRIICGDVLEVLSQIPSDSIDLGITSPPYNKKERYGGWLVDKVIYKGAKDKMNEEEYQNWQIEVLNELYRITKDGGSFFYNHKVRYEDGKMLHPVSWLLKTKWNVWQEIIWYRKIAGNIRGWRFWQVEERIYWLVKGKPKELKPEHAKFTSVWEIRPESGHKEHPAVFPIELPARIIYSILDQKGIVIDPFCGTGTTCVAAKLLGHDYIGIDISPEYVDYATKRIENTEKEKLKVISEISQHSVELTFEERKSIGLWDKKIKK | NA | NA | This protein sequence was obtained from a pMAG before dereplication during the study, so the genome is not present in the final pMAG/SAG set. The sequence showed 98.2% similarity to P-MAGv0.4_1st_NSGRY_Circle_017_413 gene, which present in the final genome set. |
| pMAG_1st_NSBR_Circle_003_1829 | CGCCTG(4,m4C) | 942 | M | II | Negative | Faintly positive | Negative | MPKKLIEVALPLEAINRQASREKSIRHGHPSTLHLWWARRPLAAARAVLFASLVDDPGEYLPPEQAERERERLFGLLERLVDWDNVKSPKEALDPKTGVIAEAQHEIAKSLARSSGQKPPTSREDRKRIEALLALAPPVLDPFAGGGTIPLEAQRLGLAAYAGDLNPVAVLINKALVDIPPRFFGLPPVNPEYRAKASQGDSFFGAQGLAWDVRYYGAWMREEAKQRLGHLYPDVEGQTVIAWIWARTVTCPNPGCRAEAPLARSFWLSRKRGKEAYIVPEVAGGKVFFRVAHEGKPPVEGTIDRRGGVCLVCGSPIPLDHVRQEGKAGRLGARLMAIVAEGQRGRNYHPPTPEHEAVAKAAVPTWKPATDLAYDPRNIWCTQYGLTTHADLFTPRQLTALATFAHLVAEAREKAYEDALKAGLPDDGVPLAQGGKGAWAYAEAVGAYLALAVDHLINRHSTLATWDASRDNVRSVFARQALPMTWDYAEANPFSEGTGAWEGMVDWVARALEALPAYPPGHARQVNALEAVNGVPTPPLISTDPPYYDNISYAALSDFFYVWLRRSLKDSYPELFRTLLVPKEEELIADPYRHGGKEAAKRRFEEGMRQVFRNLRAKAHPDYPLSLYYAFKQQEMEGEEEEEEAPKVASTGWETFLQGLVDEGFQITATWPMRTELANRPRGQGANALASSIVLVCRPRPEDAPVATRQEFLRALRRELPEALRKLTQGNVPPVDLAQSAIGPGMAVFSRYKAVLEPDGRPLSVREALTLINQVLDEFLAEEEAELDPDTRFAIAWYEQYGYGEGPYGDAETLAKAKNVAVAGLMEGGILLAKGGKVRLFRPEEYPKDWNPSADKRPSAWEAAHHLIRLLASEGESAAASLLAQLPGRLAEGARALAYRLYQVADRKGRAEDAYAFNLLAGSYGHLAVEAAKRREVQERLL | CGCCTGCAGG | EcoRV (linearize), NarI, SbfI | This protein sequence was obtained from a pMAG before dereplication during the study, so the genome is not present in the final pMAG/SAG set. The sequence showed 96.1% similarity to P-MAGv0.4_1st_NSGRN_Circle_006_1479 gene, which present in the final genome set. |
| pMAG_1st_NSBR_Circle_001_1524 | CTCGAG(1,m4C) | 937 | M | II | Negative | Negative | NA | MPKKLIEVALPLEAINEEASREKSIRHGHPSTLHLWWARRPLATARAVLFASLVDDPGEHLPEDRAAKERERLFALLERLVNWDEAKNPDSPVLWEARYEMAKALARSQGEPPPPREDREAVLALLEKAPPVLDPFAGGGTIPLEAQRLGLRAYASDLNPVAVLLNKALVEVPPLFADLPPVNPEYRSKRQPTDRFPRAKGLAEDVRFYGRWMREEAWRRIGAFYPALEGKTVIAWLWARAVACPNPACRAEAPLVRSFWLSKKRGKEAYVVPEVEGGRVVFRVKTGAGAPPREGTVNRRGAQCLVCGTPIPLEHVRKEGQAGHMGARLMAIVTEEAGGRGYHAPDPEHEVLAQDILPTWKPDFEFAKNSRHMTPWVYGLGRFSDLFTPRQLLALATFSDLVAQARERVYQDALEAGLPEDPTPLAQGGRGAWAYAEAVGVYLAMAVDRLADYHSALCSWHTGRDTIRNTFSRQALPMTWDFAEANPFSESTGSWEGMLEWVASALETLPAHPLGQARQVNAVEAVNGVPAPPLISTDPPYYDNVPYADLSDFFYVWLRRALRDTYPDLFRTLLTPKEEELIADPHRHGGKEEARRHFEEGMRRVFRHLRERAHPDYPLTLYYAFKQQEVEEDEEGETEAVASTGWETFLQGLVEEGFQVVATWPMRTELQNRPRGQGSNALASSIVLVCRPRPEDAPRATRQDFLRALRAELPRALRDLTRGSIPPVDLAQSAIGPGMAVFSRYSAVVEPDGRPLSVREALALINQVLDEFLAEEEAELDAPSRFALAWYEQYAYGEGPYGDAETLAKAKNVAVSALEEGGILLARGGKVRLFRPEEYPEDWDPKADRRLSAWEAAHHLIRLLERAGEGAAARVLAQLPPALAEGARALAYRLYQMAERKGRAEDAQSFNLLAKSYGHLAVEAARARGAVQEGLFE | ATGCAT | NarI (linearize), PstI, NsiI, BsgI | |
| pMAG_1st_NSBR_Circle_001_2404 | CTCGAG(1,m4C) | 312 | M | II | Negative | NA | NA | MRYSPVPPAQGAGEARGPSSGASYTLHIGDAREVLASFPEASVHLVVTSPPYWTLKRYEEVPGQLGHIEDYEAFLDELDRVWREVFRLLVPGGRLVVVVGDVAVARRRFGRHLVFPLHADIQVRCRKLGFDNLNPILWHKKTNASLEVEGRGVFLGKPYEPGAIIKTEVEYVLMQRKPGGYRNPTPEQRERSRLPKEDFHRFFRQIWDDIPGESTKDHPAPFPLELAERLVRMFSFVGDVVLDPFAGTGTTLIAAARWGRRALGVELVPGYAALARERFAREVPGEVLEVRGLGELETLSFTFHVQREGRYL | NA | NA | |
| pMAG_1st_NSGRY_Circle_002_1652 | GCHGC(2,m4C) | 327 | M | II | Negative | NA | NA | MTGKIRYVDLFSGAGGFSLGFHLTGYFEPALAVDNDEAAAKTFKANFPGALVLAEDIKDVSSRLIADTLGEEVDLVIGSPPCEPFTGANPRRQRNPVDRLYVDPMGQLTLHFIRIVGYLRPRVFVMENVPAIMEDGLREALIYEFRRAGYHRVFFNMLRAEEYGTPSRRRRVFVSNIPINPPRERKKVSVADALRGLPPPGRGWIPNHDPPPPLSSRKMKRAGRLGWGDAMIYYQGAERLLPNLIKLNPYDTAPTVLGSSRFIHPFENRLLTVREQARLMGFTDSFVFYGGRDEAYNMVGEAVPPPLAKAIAHEVYRYLAEEIKTLG | NA | NA | |
| pMAG_1st_NSBL_Circle_001_170 | TCGA(2,m4C) | 288 | M | II | Negative | NA | NA | MEVELNKVIVGDALTVLKTLPDEFVDVTVTSPPYNKGERDKGWLVDRIIYDAARDCKDEAEYQAEQIAVLNEIYRITKPGGSLFYNHKIRWIRGRLIHPYEWVSRSQWILRQEIIWHRKIAANLRGWRFWQVDERIYWLYKPRYEGDTVGEELNSRHALLTSVWEIPPECDERHPNPFPIELPTRCIFSVLDGRTGVVFDPYCGIGTTLVAAKLLGCDYLGIEISENYVAIARKRLENAEAERSRVLAELALHVVRKSFEERKKRGEWVGRFRKQLIKQGVLPFEGNE | NA | NA | |
| pMAG_1st_NSBL_Circle_003_539 | TGCA(4,m6A), not CTGCAG(5,m6A) | 413 | M | II | Negative | NA | NA | MEQPAAGTGEYHAETLRYMESTPMEYRVSMGQFFTPRSLREELLAHVPRLRGPRVLDPACGTGEFLLSAREYFVEPELYCWEIDGELAEVARRVVPEARVEVVDSLRKPFREEFDVVLGNPPYFEFKPDPEIRERYGEVVWGRVNVYALFVYLGIRLLKPRGYLAYVVSSSMNNGAYFRKLRDYIVRNCDIVYLRVIDDPYVFKEVNHTFQLLVLRKGRNTGRYVFRKGNTIVFSERAEELRRVFESSKTLEELGYRVMTGRVVWNQHRDKLTDDPNEGILLVWAHNIKHGRLVLNNRDRPQYIRWPLDRADRGPAIVVRRVVGHPRRLRLEAALVPPGTVFVAENHVNVVYPPPGATPEEMEEIVEQLNSEETSRILSAIIGNTQVSRRELGSLVPLAIATAGAPSASSRNL | NA | NA | |
| P-MAGv0.4_1st_NSBL_Circle_006_879 | TGCA(4,m6A), not CTGCAG(5,m6A) | 422 | M | II | Negative | NA | NA | MIKSGELHWETVKYIMRADPGKRASLGQFFTPRALREELLSRLPRLHRPKVLDPACGTGEFLLSAREYFVEPELYCWEVDGELAEIARRIVPEAKIEVVDSLRKPFREEFDVVIGNPPYFEFKPDPEIVERYREVIWGRVNVYALFVYLGVKLLKPGGYLAYVVSSSMNNGAYFRKLREFLVRNCDIVYLKRIDDPYVFSDPYYKVNHTFQLLVLRKGENTGRYVFKHRGIMIFTEEYEYLRRVFEESVTLKDLGYRVLTGRVVWNQNRDKLTDNPKEGILLIWAHNIKRGKLILNNRERPQYIKWPIDRADKGPAIVVTRVVGHPGKASLEAALVPPGVIFVAENHVNVVYPPPHATLEEMEEIVEQLNSEEASKIVKKITGNTQISKTELENLIPLRLRSRSVLKSTKHNLLEYLRQTR | NA | NA | |
| pMAG_1st_NSGRY_Circle_003_930\|LOCUS_09220 | TGCA(4,m6A), not CTGCAG(5,m6A) | 460 | M | II | Negative | NA | NA | MRSERPRSRPELGSGAAWLCSYIPGAVLTRGTSTQLGVLHYCPRMMAVLVDEYHPETVRYLRESDPGHRRRLGQFFTPRRLREELLSKIPRLIEPRVLDPACGTGEFLLTAKQYFVEPELHCWEIDPRLAEIARGVVPEARVEVVDALLKPFREEFDAVIGNPPYFEFKPSGEIEARFREIVYGRPNIYAFFIYLGLRVLKPGGYLAYVVSSSMNNGAYFKKLREFIVRCADIVHLKTLRDQVLFEGVNHTFQLLVLRKTGNTGRYVFEHSGLMIFSEYSDYLRKAFKGSLTLRELGYRVKTGRVVWNQHRDKLTDDPSQGILLVWAHNIKRGRLVLGNRPNKPQYIKWPPEKADEGPAIVATRIVGHPGSASIDAALVPPGTVFVAENHVNVIYPPPTASLRDVEEVVRQLNSPETRRIIRAITGNTQISKTELERLLPVKPVTPGHRVDQVPHRPSLL | NA | NA | |

### Supplementary Data

**Data S1. Summary of MAGs and SAGs.**

**Data S2. Methylated motifs detected in MAGs and SAGs.**

**Data S3. RM system genes predicted in MAGs and SAGs.**

**Data S4. Methylation ratio of methylated motifs.**
